# Chromosome-level reference genomes for two strains of *Caenorhabditis briggsae*: an improved platform for comparative genomics

**DOI:** 10.1101/2021.12.01.470807

**Authors:** Lewis Stevens, Nicolas D. Moya, Robyn E. Tanny, Sophia B. Gibson, Alan Tracey, Huimin Na, Ye Zhan, Rojin Chitrakar, Job Dekker, Albertha J.M. Walhout, L. Ryan Baugh, Erik C. Andersen

**Author notes:** **Corresponding author:** Erik C. Andersen, Department of Molecular Biosciences, Northwestern University, 4619 Silverman Hall, 2205 Tech Drive, Evanston, IL 60208. Tree of Life, Wellcome Sanger Institute, Cambridge, UK.

## Abstract

The publication of the *Caenorhabditis briggsae* reference genome in 2003 enabled the first comparative genomics studies between *C. elegans* and *C. briggsae*, shedding light on the evolution of genome content and structure in the *Caenorhabditis* genus. However, despite being widely used, the currently available *C. briggsae* reference genome is substantially less complete and structurally accurate than the *C. elegans* reference genome. Here, we used high-coverage Oxford Nanopore long-read and chromosome conformation capture data to generate chromosomally resolved reference genomes for two *C. briggsae* strains: QX1410, a new reference strain closely related to the laboratory AF16 strain, and VX34, a highly divergent strain isolated in China. We also sequenced 99 recombinant inbred lines (RILs) generated from reciprocal crosses between QX1410 and VX34 to create a recombination map and identify chromosomal domains. Additionally, we used both short- and long-read RNA sequencing (RNA-seq) data to generate high-quality gene annotations. By comparing these new reference genomes to the current reference, we reveal that hyper-divergent haplotypes cover large portions of the *C. briggsae* genome, similar to recent reports in *C. elegans* and *C. tropicalis*. We also show that the genomes of selfing *Caenorhabditis* species have undergone more rearrangement than their outcrossing relatives, which has biased previous estimates of rearrangement rate in *Caenorhabditis*. These new genomes provide a substantially improved platform for comparative genomics in *Caenorhabditis* and narrow the gap between the quality of genomic resources available for *C. elegans* and *C. briggsae*.

## Introduction

Since its introduction as a model for animal development in the 1950s, the free-living nematode *Caenorhabditis elegans* has become one of the key model organisms in biology. However, before eventually settling on *C. elegans*, Sydney Brenner had initially proposed a closely related species, *Caenorhabditis briggsae*, for study in the laboratory (Edgar and Wood 1977; Riddle et al. 2011). Although *C. briggsae* has received far less attention than its close relative, it is now widely used as a satellite model organism and has played a key role in our understanding of *Caenorhabditis* evolution, development, and genetics (Baird and Chamberlin 2006). *C. elegans* and *C. briggsae* share many features of their biology: they are nearly morphologically indistinguishable; both species reproduce predominantly by self-fertilization; and both are found globally, often in the same local habitats (Nigon and Dougherty 1949; Cutter et al. 2006; Barrière and Félix 2005; Dolgin et al. 2007; Félix and Duveau 2012; Thomas et al. 2015; Crombie et al. 2019). However, it is their differences that have helped shed light on several key biological pathways. For example, *C. elegans* and *C. briggsae* independently evolved from outcrossing species to reproduce as selfing hermaphrodites (which are essentially females capable of producing sperm) and have done so using distinct genetic mechanisms (Cho et al. 2004; Kiontke et al. 2011; Nayak et al. 2005; Hill et al. 2006).

In 2003, five years after the *C. elegans* reference genome was published (*C. elegans* Sequencing Consortium 1998), a reference genome for the laboratory strain of *C. briggsae*, AF16, was generated using Sanger-based shotgun sequencing and a physical map generated using fosmids and bacterial artificial chromosomes (BACs) (Stein et al. 2003). This genome was subsequently resolved into chromosomes using a genetic map constructed by sequencing lines from inter-strain crosses (Hillier et al. 2007; Ross et al. 2011). The availability of a high-quality reference genome for *C. briggsae* enabled the first comparative genomics studies between *C. elegans* and *C. briggsae*, revealing that their genomes have diverged substantially since they last shared a common ancestor, and have undergone a strikingly high rate of intrachromosomal rearrangement (Coghlan and Wolfe 2002). Despite these differences, the higher-order structure of the *C. elegans* genome is largely conserved in *C. briggsae* and surprisingly few genes have moved between chromosomes (Hillier et al. 2007). The reference genome also provided a foundation for population genetic and genomic studies, revealing that *C. briggsae* harbors higher levels of genetic diversity than *C. elegans*, largely driven by the existence of distinct phylogeographic groups and makes *C. briggsae* a more suitable model for studying gene flow and speciation (Thomas et al. 2015). Nearly 20 years later, the *C. briggsae* reference genome remains one the highest quality *Caenorhabditis* genomes currently available (Harris et al. 2020). However, it still lags behind the *C. elegans* genome in terms of accuracy and completeness, containing thousands of gaps that are estimated to comprise several megabases of sequence along with numerous regions that have been found to be misoriented (Ren et al. 2018).

Since the *C. briggsae* reference genome was published, DNA sequencing technology has advanced at an exponential rate. In recent years, long-read sequencing technologies, namely those offered by Pacific Biosciences (PacBio) and Oxford Nanopore Technologies (ONT), have revolutionized genome assembly, and it is now relatively straightforward to generate highly contiguous genome assemblies (Rhie et al. 2021). Moreover, high-throughput mapping technologies, such as chromosome-conformation capture (Hi-C), are now routinely being used to construct fully chromosomally resolved reference genomes that far surpass the quality of their predecessors (Rhie et al. 2021). Similarly, it is now possible to generate long-read RNA sequencing data to assemble full-length transcripts and substantially improve the quality of automated gene annotations. Substantially improved reference genomes generated using these technologies have recently been published for several nematode species, including for *Caenorhabditis tropicalis* and *Caenorhabditis remanei* (Noble et al. 2021; Teterina et al. 2020), *Oscheius tipulae* (Gonzalez de la Rosa et al. 2021), and the ruminant parasite *Haemonchus contortus* (Doyle et al. 2020). *These technologies have even been used to identify and extend multiple repetitive regions that are collapsed in the C. elegans* N2 reference genome (Tyson et al. 2018; Yoshimura et al. 2019), widely regarded as one of the highest quality eukaryotic reference genomes available.

Here, we use the Oxford Nanopore PromethION platform and Hi-C to sequence the genomes of two *C. briggsae* strains, QX1410, a new reference strain closely related to AF16, and VX34, a highly divergent strain. QX1410 was chosen instead of improving the AF16 genome because the fidelity of the AF16 strain is no longer certain after its long time in the laboratory. We use these data to generate two high-quality reference genomes that have biological completeness scores equal to that of the current *C. elegans* reference genome, and they have substantially fewer gaps and unplaced sequence than the existing *C. briggsae* AF16 reference genome. We also use both long- and short-read RNA sequencing (RNA-seq) data to generate high-quality, automated gene annotations that are comparable in quality to the curated AF16 annotations. We also generated and sequenced a panel of recombinant inbred lines from reciprocal crosses between QX1410 and VX34 to characterize the recombination landscape of the *C. briggsae* genome. Consistent with previous reports in *C. elegans* and *C. tropicalis*, we find that hyper-divergent haplotypes cover large portions of the *C. briggsae* genome. We also revisit one of the first comparative genomics analyses performed between *C. briggsae* and *C. elegans* and reveal that the genomes of selfing *Caenorhabditis* species have undergone more rearrangement than their outcrossing relatives, which has biased previous estimates of rearrangement rate in *Caenorhabditis*. These new resources will provide a substantially improved foundation for genomic analyses in this important satellite model organism.

## Results

### High-quality reference genomes for two *C. briggsae* strains

We sought to generate a high-quality reference genome for QX1410, a new reference strain for *C. briggsae* isolated from Saint Lucia that is closely related to AF16, the current reference strain (Thomas et al. 2015). To facilitate comparative analyses, we also sequenced the genome of a highly divergent strain, VX34, isolated from China. We generated high-coverage long-read data for each strain (read length N50s of 23.6 kb and 23.5 kb, respectively; coverages of 219x and 508x, respectively) using the Oxford Nanopore PromethION platform and sequenced chromosome-conformation capture (Hi-C) libraries for each strain to high coverage (173x and 168x, respectively) using Illumina technology. We assembled the long PromethION reads independently for each strain using several tools and chose the most contiguous assemblies, both of which comprised several contigs that represented complete chromosomes (Fig. S1). We corrected sequencing errors in the contigs, and these polished contigs were then scaffolded into complete chromosomes using the Hi-C data (Figure S2). Each reference genome was manually curated by inspecting coverage of the long-reads and by assessing congruence with the Hi-C contact maps.

The resulting reference genomes for QX1410 and VX34 span 106.2 Mb and 107.0 Mb, respectively, similar in span to the existing AF16 reference (Table 1). Both comprise six scaffolds representing the six chromosomes (I-V, X), a complete mitochondrial genome, and have no unplaced sequence. The QX1410 and VX34 assemblies have high base-pair accuracy, with consensus quality value (QV) scores of 45.6 (which corresponds to one error in 36.2 kb) and 44.4 (one error in 27.5 kb), respectively. BUSCO completeness is equal to that of the current *C. elegans* N2 reference genome (99.4%), compared with 98.8% for the existing *C. briggsae* AF16 reference (Table 1). The new reference genomes are substantially more contiguous than AF16, with contig N50s of 14.7 Mb and 16.1 Mb respectively, compared with 47 kb. Only seven gaps remain in the QX1410 reference genome (one in chromosome I, one in chromosome II, and five in the X chromosome) and three gaps in the VX34 reference genome (two in chromosome in V and one in the X chromosome) compared with 4,706 gaps in the existing AF16 reference genome. In contrast to the chromosomal scaffolds in AF16, all of which lack the nematode telomeric repeat sequence ([TTAGGC]n) at their ends, a majority of the chromosomes in our reference genomes end in telomeric repeat sequences (Fig. S3).

**Table 1:**
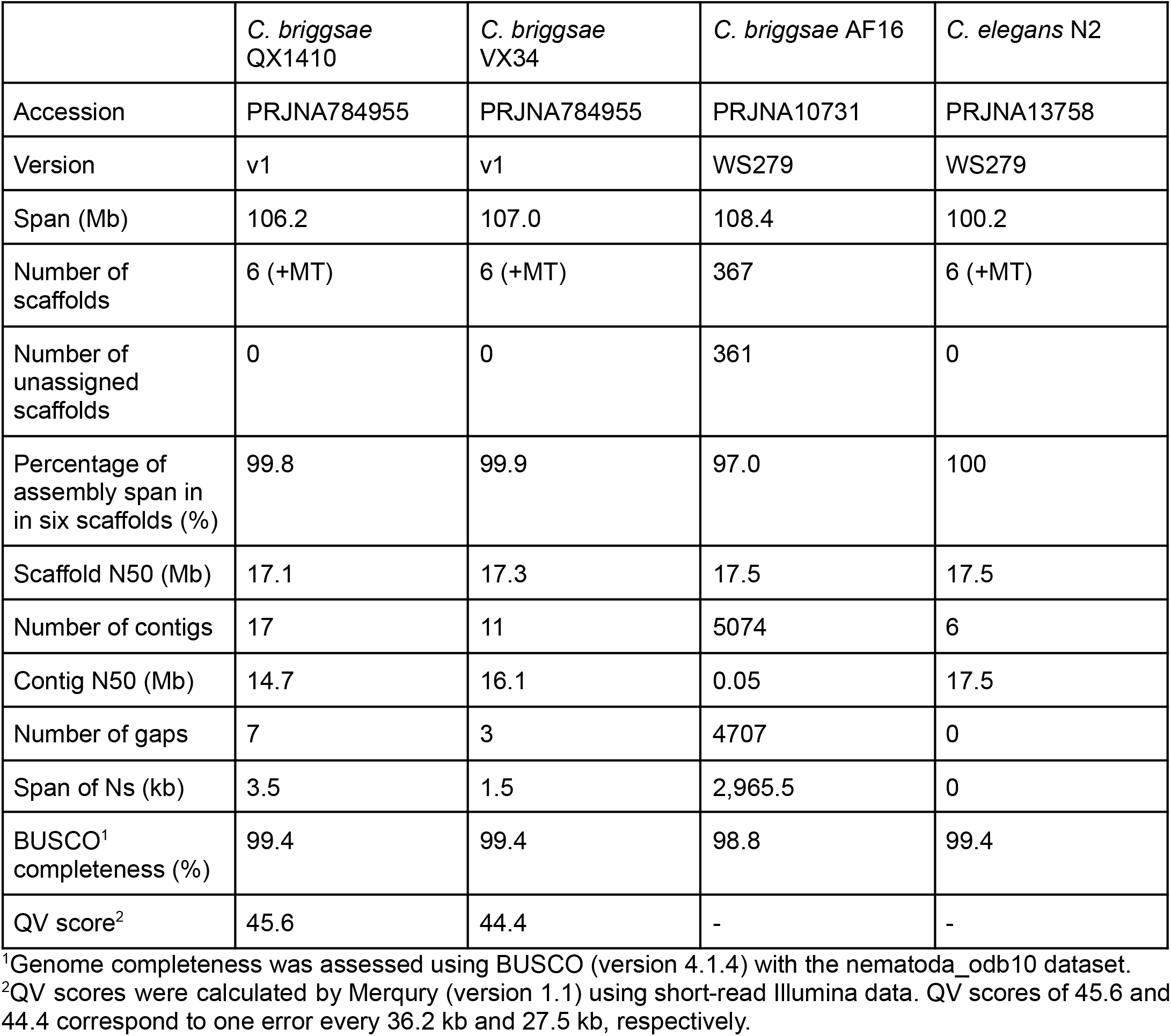
Reference genome metrics.

|  | *C. briggsae*<br>QX1410 | *C. briggsae*<br>VX34 | *C. briggsae* AF16 | *C. elegans* N2 |
| --- | --- | --- | --- | --- |
| Accession | PRJNA784955 | PRJNA784955 | PRJNA10731 | PRJNA13758 |
| Version | v1 | v1 | WS279 | WS279 |
| Span (Mb) | 106.2 | 107.0 | 108.4 | 100.2 |
| Number of scaffolds | 6 (+MT) | 6 (+MT) | 367 | 6 (+MT) |
| Number of unassigned scaffolds | 0 | 0 | 361 | 0 |
| Percentage of assembly span in in six scaffolds (%) | 99.8 | 99.9 | 97.0 | 100 |
| Scaffold N50 (Mb) | 17.1 | 17.3 | 17.5 | 17.5 |
| Number of contigs | 17 | 11 | 5074 | 6 |
| Contig N50 (Mb) | 14.7 | 16.1 | 0.05 | 17.5 |
| Number of gaps | 7 | 3 | 4707 | 0 |
| Span of Ns (kb) | 3.5 | 1.5 | 2,965.5 | 0 |
| BUSCO<sup>1</sup> completeness (%) | 99.4 | 99.4 | 98.8 | 99.4 |
| QV score<sup>2</sup> | 45.6 | 44.4 | - | - |

<sup>1</sup>Genome completeness was assessed using BUSCO (version 4.1.4) with the nematoda\_odb10 dataset.
<sup>2</sup>QV scores were calculated by Merqury (version 1.1) using short-read Illumina data. QV scores of 45.6 and 44.4 correspond to one error every 36.2 kb and 27.5 kb, respectively.

Despite having high-coverage long-read data, manual curation revealed that the subtelomeric regions, which are known to be highly repetitive in *C. elegans* (Kim et al. 2019), are unresolved in five of the 12 ends of the QX1410 reference genome. One of these is the left-end of chromosome V (VL), which ends in nine tandemly repeated ∼7.5 kb ribosomal DNA (rDNA) cistron units (Figure S4A). In C. elegans, the rDNA cluster consists of approximately 55 repeated units and sits adjacent to the telomeric repeat sequence on chromosome I (Ellis et al. 1986; C. elegans Sequencing Consortium 1998). In QX1410, the average coverage within this repeated region is 943x, approximately 6.5-times higher than the chromosome-wide average (∼146x), suggesting that this region is collapsed in our assembly and that the true number of rDNA repeats in QX1410 is approximately 58 (Figure S4B). Another unresolved region is the right-end of the X chromosome (XR), which ends in a large ∼65 kb tandem repeat (Figure S5A) that presumably prevented the genome assembler from extending into the telomeric repeat sequence. Two of the remaining unresolved chromosome ends, IL and IVL, end in unique sequences that are punctuated with blocks of kmers that contain the nematode telomeric sequence. For example, in the IVL subtelomere, we find multiple blocks of a repeating 19-mer (TTAGGCTTAGGCTTCCCGC) interspersed with a unique sequence (Figure S5B; Figure S6A). Blocks of the same sequence are also found in the IVR subtelomere (Figure S6A). In the IL subtelomere, we find a block of a repeating 14-mer (TAAGCCTAAGCCTC) with blocks of the same sequence also found on the IIR and XL (Figure S6A). Similar blocks of both kmers exist in the subtelomeric regions of the VX34 genome, but they are found at subtelomeres of different chromosomes (Figure S6B).

The AF16 reference genome has previously been reported to contain multiple mis-scaffolded regions (Ren et al. 2018). Although efforts have been made to correct these regions, these changes are not yet reflected in the current version of the reference available via public databases (Harris et al. 2020). To assess collinearity between the three *C. briggsae* reference genomes, we aligned AF16 and VX34 to the QX1410 reference genome. Consistent with previous reports, we found numerous regions in all six chromosomes that are inverted in AF16 relative to QX1410 (Figure 1C). By contrast, and despite being far more divergent from QX1410 than AF16 and being assembled independently, the VX34 genome is highly collinear with the QX1410 reference genome (Figure 1B), suggesting that the AF16 genome is not accurate. To quantify this relationship, we called structural variants in the reference genomes of AF16 and VX34 using QX1410 as the reference and focused on large inversions (≥ 50 kb in length). Relative to QX1410, AF16 has 47 inversions (average size of 167 kb) and VX34 has six (average 83 kb). Of those inversions above 100 kb, AF16 has 31 while VX34 has just one. Importantly, the QX1410 and VX34 reference genomes were generated by scaffolding contigs that spanned multiple megabases (contig N50s of 14.7 Mb and 16.1 Mb, respectively), meaning that all of the inversions called in VX34 fall within contiguous sequence and are thus likely to represent real structural variation. In contrast, the AF16 reference genome was scaffolded using a highly fragmented assembly (contig N50 of 47 kb), meaning many of the observed variants are likely to be scaffolding errors. In summary, the new reference genomes we have generated for QX1410 and VX34 are substantially more contiguous, complete, and structurally correct than the current AF16 reference genome.

**Figure 1:**
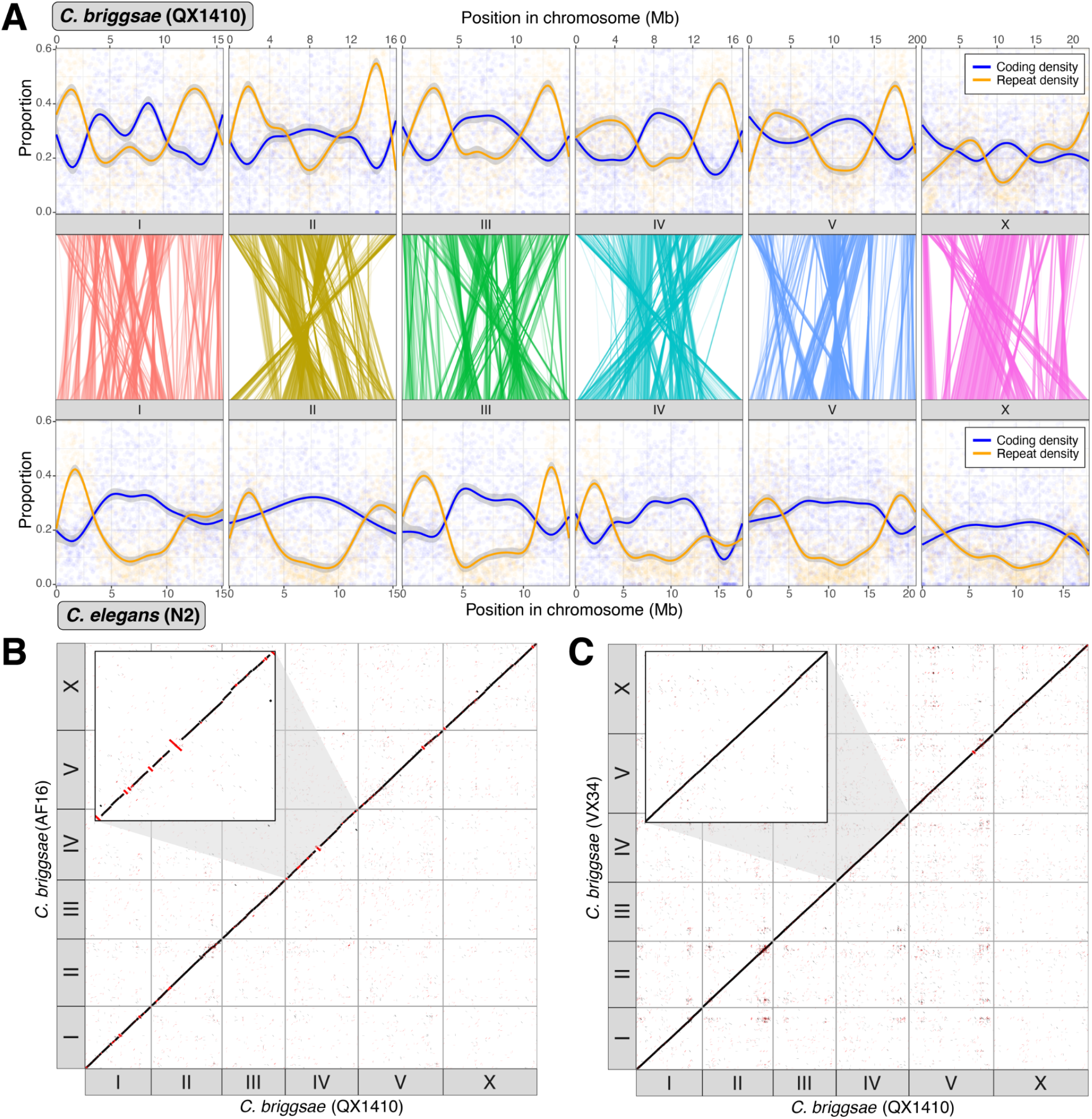
High-quality reference genomes for two *C. briggsae* strains. A) Comparison between the *C. briggsae* QX1410 and *C. elegans* N2 reference genomes. Repeat density and protein-coding gene density per 10 kb windows are shown. Repeats were identified *de novo* using RepeatModeler2. Solid lines represent LOESS smoothing functions fitted to the data. Relative positions of 9,461 one-to-one orthologs are shown as lines joining the two density plots. B) Whole-genome alignment of AF16 to the QX1410 reference genome generated using nucmer. Alignments shorter than 1 kbp are not shown. Alignments in the reverse orientation are highlighted in red. Inset: chromosome IV showing multiple regions between AF16 and QX1410 that are in different orientations. C) Whole-genome alignment of VX34 to the QX1410 reference genome generated using nucmer. Alignments shorter than 1 kbp are not shown. Alignments in the reverse orientation are highlighted in red. Inset: the same chromosome IV region as in (B) showing a largely collinear alignment.

### High-quality protein-coding gene annotations using long- and short-read RNA-seq

We sought to develop a computational pipeline that leverages long- and short-read RNA-seq reads to generate high-quality protein-coding gene annotations for the QX1410 and VX34 reference genomes. We collected RNA from mixed-stage, male-enriched, and starved cultures to maximize transcript detection. To allow us to use the highly curated *C. elegans* N2 reference annotation (PRJNA13758) as a truth set, we sequenced the transcriptome of *C. elegans* strain PD1074 (a recent clone of N2) using the Pacific Biosciences (PacBio) Single-Molecule Real-Time (SMRT) platform and benchmarked several transcriptome assembly and gene prediction tools. We refined the PacBio long-reads into 55,936 high-quality transcripts and generated gene models by predicting open reading frames (ORFs) in the high-quality transcripts. Additionally, we generated protein-coding gene predictions from short RNA-seq read alignments. Based on our N2 benchmark, we merged the best transcriptome assembly and gene prediction models into a single, non-redundant gene set. The BUSCO completeness of our gene set was 99.1%, only slightly lower than the completeness of the existing *C. elegans* N2 reference gene set (99.2%). Merging the short- and long-read based gene models led to improvements in exon, intron, and transcript sensitivity relative to either approach alone (Table S1). Our final annotation has correct predictions for 74.7% and 86.1% of all *C. elegans* exons and introns, respectively. We correctly predicted 43.1% of all *C. elegans* transcripts, with at least one correctly predicted transcript in 68.2% of the *C. elegans* genes.

Using this same process, we sequenced the transcriptomes of both QX1410 and VX34 strains using PacBio SMRT and Illumina platforms and employed our pipeline to generate high-quality gene models for both *C. briggsae* strains. We predicted 21,514 and 20,023 protein-coding genes in the QX1410 and VX34 genomes, respectively, similar to the number of genes predicted for AF16 (20,821). BUSCO completeness of the QX1410 and VX34 annotations is equal to the current AF16 (WS280) annotation (99.2%), which has undergone extensive manual curation. We also assessed gene annotation quality by comparing the length of each gene in our gene sets to their corresponding orthologs in *C. elegans* (Fig. S7). We performed the same protein length analysis for the current *C. briggsae* AF16 reference annotation (WS280) and a version of the AF16 annotation composed of automated predictions only (WS255). Our QX1410 and VX34 annotations show substantial improvements in protein-length accuracy relative to the uncurated AF16 WS255 release (Table S2). However, compared to the AF16 WS280 release, we find that our annotations have fewer matches that are identical in length or within 5% of the length of the orthologous *C. elegans* protein (Table S2). Manual curation is needed to further improve the QX1410 and VX34 gene models to bring these annotations to protein-length accuracies comparable to the manually curated AF16 WS280 release.

### Recombination in the *C. briggsae* recombinant lines

To characterize the recombination landscape between QX1410 and VX34, we generated a panel of 99 F2 recombinant inbred lines (RILs) from reciprocal crosses between the two strains and genotyped 2,981 single nucleotide markers in all RILs. After removing markers with distorted segregation patterns and markers with large deviations in frequency relative to neighbors (Figure S8, Table S3), the remaining 2,828 markers were used to estimate a genetic map for *C. briggsae*. The estimated genetic map has a length of 508.8 cM and spans 97.7% of the QX1410 genome physical length. The size of this genetic map is similar to that of previous estimations using an F2 RIL scheme (588.1 cM) (Hillier et al. 2007) and approximately 55% of the size of previous estimations using an advanced intercross (AI) scheme (928.6 cM) (Ross et al. 2011). The reduction in size relative to this previous map can be explained, at least in part, by the reduced number of recombination breakpoints expected in RILs relative to recombinant lines from advanced intercross (Rockman and Kruglyak 2008, 2009).

As in *C. elegans*, the genetic maps for the six *C. briggsae* chromosomes have distinct arms and centers that show detectable changes in recombination rate (Hillier et al. 2007; Rockman and Kruglyak 2009; Ross et al. 2011). We generated Marey maps to show the genetic position as a function of the physical position and used segmented linear regression to identify arm-center boundaries and estimate the rate of recombination in each domain (Figure 2A, Table 2). Most of our domain boundaries are in agreement with the span and physical position of previously defined domains (Ross et al. 2011). Although the recombination rate is relatively constant within each domain, we observed small segments in chromosomal arms where recombination rate abruptly approached zero. This pattern can be explained by sampling error, where the limited number of recombination events that occur across the 99 RILs could lead to several genomic markers with skewed recombination fractions. With the exception of IIL, the recombination rate at chromosome ends sharply decreased, expected for regions approaching chromosome arm-tip boundaries. These RILs clearly resolve tip domains at IIIR, IVL, IVR, and XL. Tip domain boundaries were manually defined.

**Table 2:**
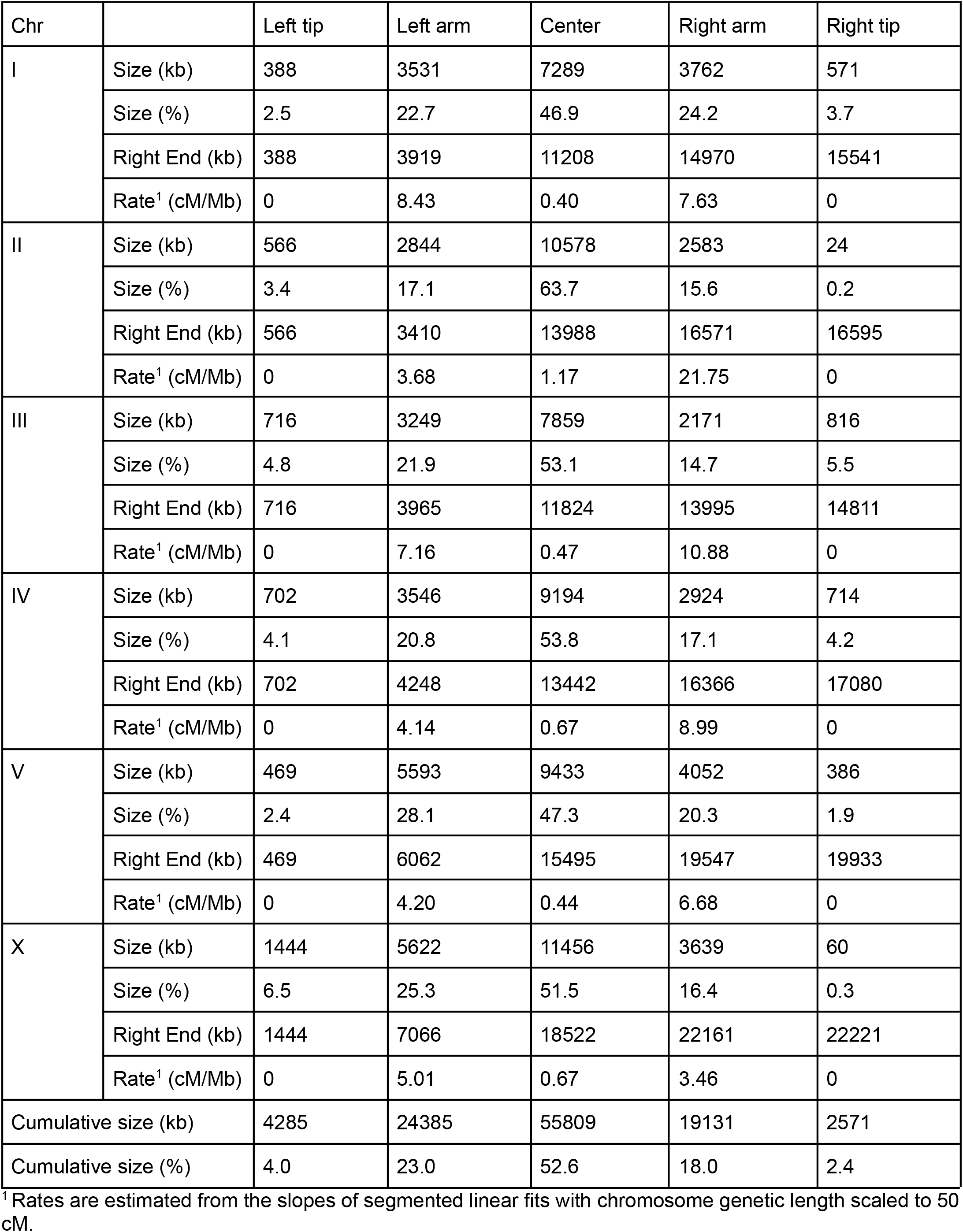
Chromosomal Domains.

| Chr |  | Left tip | Left arm | Center | Right arm | Right tip |
| --- | --- | --- | --- | --- | --- | --- |
| I | Size (kb) | 388 | 3531 | 7289 | 3762 | 571 |
|  | Size (%) | 2.5 | 22.7 | 46.9 | 24.2 | 3.7 |
|  | Right End (kb) | 388 | 3919 | 11208 | 14970 | 15541 |
|  | Rate<sup>1</sup> (cM/Mb) | 0 | 8.43 | 0.40 | 7.63 | 0 |
| II | Size (kb) | 566 | 2844 | 10578 | 2583 | 24 |
|  | Size (%) | 3.4 | 17.1 | 63.7 | 15.6 | 0.2 |
|  | Right End (kb) | 566 | 3410 | 13988 | 16571 | 16595 |
|  | Rate<sup>1</sup> (cM/Mb) | 0 | 3.68 | 1.17 | 21.75 | 0 |
| III | Size (kb) | 716 | 3249 | 7859 | 2171 | 816 |
|  | Size (%) | 4.8 | 21.9 | 53.1 | 14.7 | 5.5 |
|  | Right End (kb) | 716 | 3965 | 11824 | 13995 | 14811 |
|  | Rate<sup>1</sup> (cM/Mb) | 0 | 7.16 | 0.47 | 10.88 | 0 |
| IV | Size (kb) | 702 | 3546 | 9194 | 2924 | 714 |
|  | Size (%) | 4.1 | 20.8 | 53.8 | 17.1 | 4.2 |
|  | Right End (kb) | 702 | 4248 | 13442 | 16366 | 17080 |
|  | Rate<sup>1</sup> (cM/Mb) | 0 | 4.14 | 0.67 | 8.99 | 0 |
| V | Size (kb) | 469 | 5593 | 9433 | 4052 | 386 |
|  | Size (%) | 2.4 | 28.1 | 47.3 | 20.3 | 1.9 |
|  | Right End (kb) | 469 | 6062 | 15495 | 19547 | 19933 |
|  | Rate<sup>1</sup> (cM/Mb) | 0 | 4.20 | 0.44 | 6.68 | 0 |
| X | Size (kb) | 1444 | 5622 | 11456 | 3639 | 60 |
|  | Size (%) | 6.5 | 25.3 | 51.5 | 16.4 | 0.3 |
|  | Right End (kb) | 1444 | 7066 | 18522 | 22161 | 22221 |
|  | Rate<sup>1</sup> (cM/Mb) | 0 | 5.01 | 0.67 | 3.46 | 0 |
| Cumulative size (kb) |  | 4285 | 24385 | 55809 | 19131 | 2571 |
| Cumulative size (%) |  | 4.0 | 23.0 | 52.6 | 18.0 | 2.4 |

<sup>1</sup> Rates are estimated from the slopes of segmented linear fits with chromosome genetic length scaled to 50 cM.

**Figure 2:**
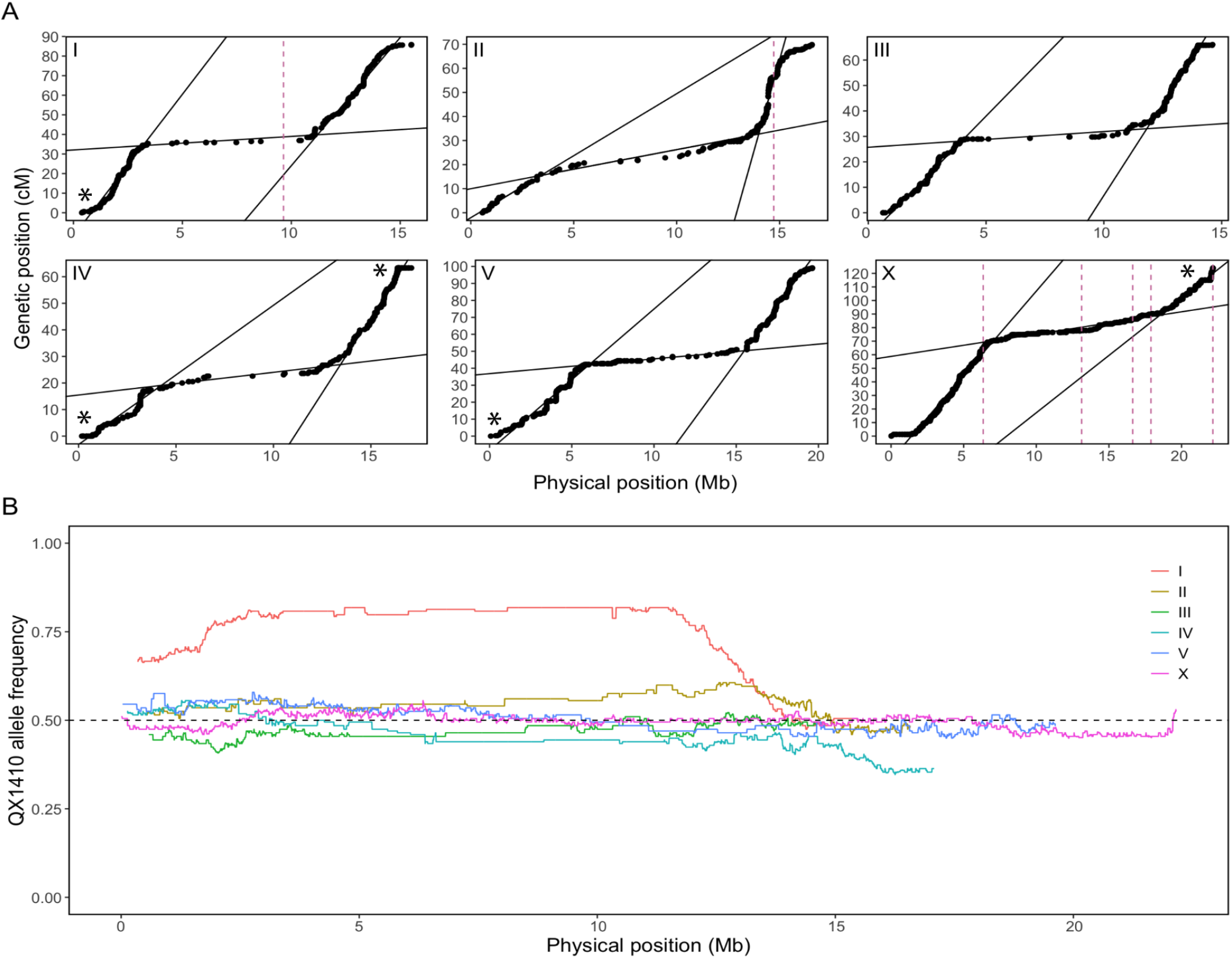
Recombination rates in the *C. briggsae* genome determined by genotyping 99 QX1410xVX34 RILs. (A) Marey maps for each chromosome in QX1410. The genetic position of each marker is shown as a function of physical position (black dots). Fits from segmented linear regressions are shown as black lines. Changepoints in the segmented linear regressions were used to estimate chromosome domain boundaries and the rate of recombination. Dashed pink lines indicate the physical position of gaps in the QX1410 genome assembly. Asterisks were added to chromosomal ends where subtelomeric regions are unresolved. (B) Frequency of the QX1410 allele as a function of physical position across every marker in each chromosome. Allele frequency was averaged using a sliding window 100 kb with a step size of 5 kb. The neutral expected frequency of 0.5 is shown as a dashed horizontal black line.

In selfing *Caenorhabditis* species, evidence suggests that selective pressure to maintain linkage between coadapted alleles might lead to incompatibilities between interbreeding populations (Seidel et al. 2008; Rockman and Kruglyak 2009; Ross et al. 2011; Noble et al. 2021). A strong departure from the neutral expectation in allele frequencies was observed in chromosomes I, II, and IV (Figure 2B). These skews might be explained by regions of incompatibility between QX1410 and VX34. Interestingly, the skew in chromosome I spans the entirety of the chromosome center. Although this skew may be the result of massive chromosome I incompatibilities between the two strains, markers in chromosome I center are sparse. Incompatibilities surrounding these sparse markers might drive an inflation in average allele frequencies across this genomic interval.

### High divergence among the *C. briggsae* reference genomes

The genomes of *C. elegans* wild isolates contain large, hyper-divergent haplotypes that comprise unique sets of genes and alleles that are highly diverged at the amino acid level (Thompson et al. 2015; Lee et al. 2021). These haplotypes are hypothesized to be remnants of genetic diversity present in the outcrossing ancestor that have been maintained by long-term balancing selection since the evolution of selfing (Thompson et al. 2015; Lee et al. 2021). Similar hyper-divergent regions have been reported in both *C. briggsae* (Lee et al. 2021) and in the related selfing species, *C. tropicalis* (Noble et al. 2021; Ben-David et al. 2021). To quantify the genome-wide divergence between the three sequenced *C. briggsae* strains, we aligned the AF16 and VX34 reference genomes to QX1410, calculated alignment identity, and called variants (SNVs and indels). Given that hyper-divergent haplotypes in *C. elegans* often show little homology to each other at the nucleotide level, we also identified alleles across all three strains using an orthology clustering approach and calculated amino acid identity.

Consistent with recent findings in *C. elegans* and *C. tropicalis* (Lee et al. 2021; Noble et al. 2021), we find that hyper-divergent haplotypes are widespread across the genomes of all three *C. briggsae* strains (Fig. 3). Despite QX1410 and AF16 belonging to the same “Tropical” genetic group (Thomas et al. 2015), large regions of their genomes are unalignable at the nucleotide level (Figure 3A). As in *C. elegans*, these regions are overrepresented in the autosomal arms and underrepresented in the autosomal centers and on the X chromosome (Lee et al. 2021). The nucleotide divergence in these regions is associated with a substantial drop in amino acid identity between alleles, with many alleles showing less than 95% identity (Figure 3A). For example, large regions of the right-arm of chromosome II are unalignable between QX1410 and AF16, and alleles across this arm have a mean amino acid identity of 98.3% (Figure 3A). Comparing QX1410 with the more divergent VX34, we find a substantial divergence in the arms of all chromosomes (Figure 3B). As in AF16, the right-arms of chromosome II and V are particularly divergent, with an mean amino acid identity of 97.4% and 96.8% between alleles, respectively (Figure 3B). Surprisingly, we also find substantial divergence on the X chromosome between QX1410 and VX34, with a mean amino acid identity of 97.8% across its length and large sections of the chromosome arms show poor or no alignment (Figure 3B).

**Figure 3:**
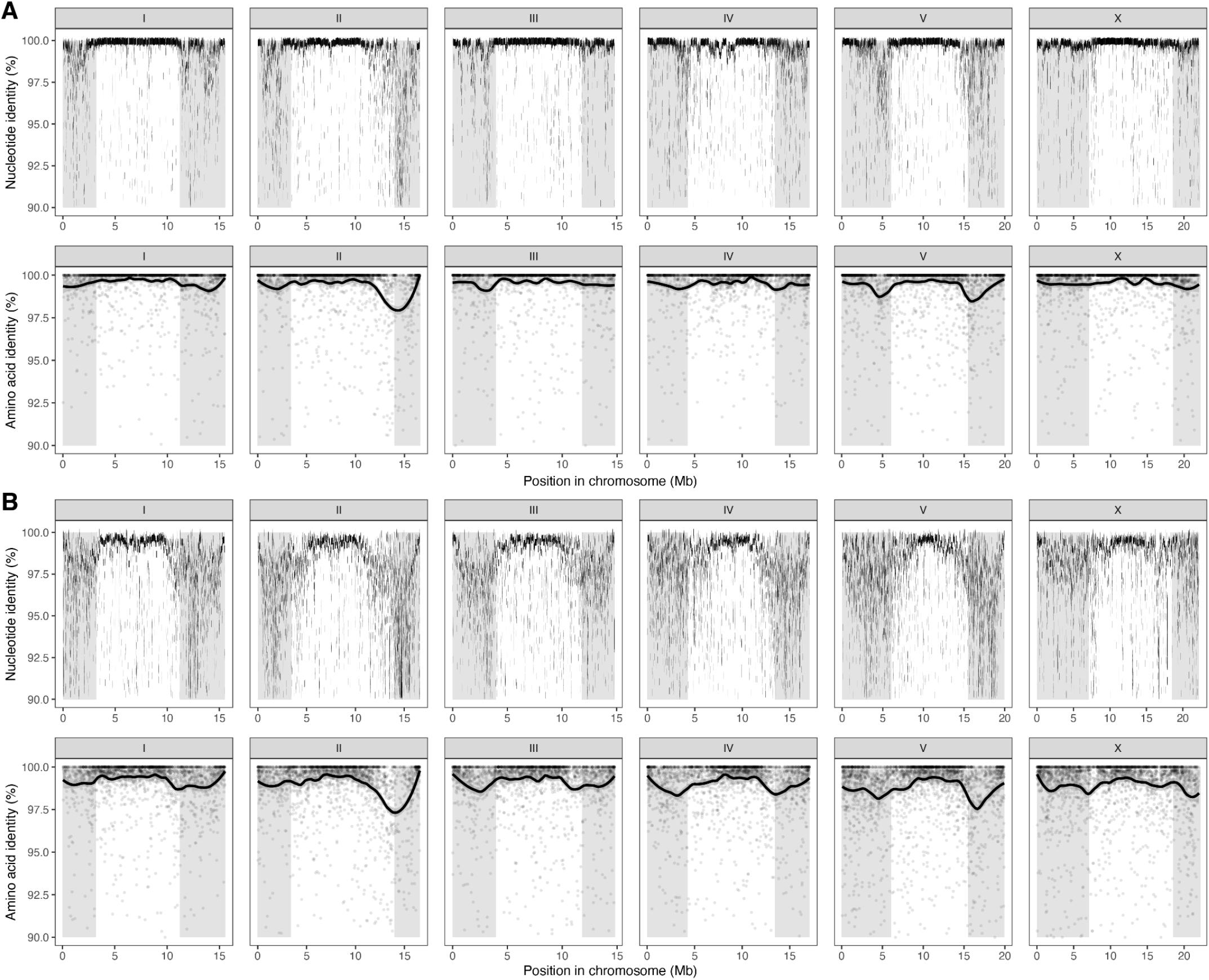
Genome-wide divergence between three *C. briggsae* strains. Genomes were aligned using nucmer and aligned regions of 1 kb or longer are shown. Conserved protein sequences were identified using OrthoFinder and aligned using MAFFT; lines represent LOESS smoothing curves fitted to the amino acid identity data. Grey shading indicates chromosome arm regions defined previously. (A) Nucleotide identity between aligned regions of QX1410 and AF16 genomes, and amino acid identity of protein sequences conserved between QX1410 and AF16. (B) Nucleotide identity between aligned regions of QX1410 and VX34 genomes, and amino acid identity of protein sequences conserved between QX1410 and VX34.

To place the level of divergence in *C. briggsae* in the context of previous work in *C. elegans*, we also compared aligned long-read assemblies for CB4856 (a divergent strain from Hawaii) and XZ1516 (the most divergent *C. elegans* strain currently known, also isolated in Hawaii) to the laboratory N2 reference genome and called variants (SNVs and indels). As expected, the divergence between the QX1410 and AF16 is the lowest of all the comparisons, with one SNV every 634 bp. The divergence between QX1410 and VX34 (average of one SNV every 135 bp) is substantially higher than the higher divergence between the *C. elegans* strains N2 and CB4856 (one SNV every 362 bp, respectively) (Figure S9A-C) and similar to that between N2 and the most divergent *C. elegans* strain XZ1516 (one SNV in 141 bp) (Figure S9D). Importantly, the distribution of divergence is qualitatively different across comparisons. Divergence is higher between VX34 and QX1410 than between N2 and XZ1516 across all chromosomes other than chromosome V, which harbors a notable excess of SNVs relative to the other autosomes in *C. elegans* (Figure S10). We also find that the X chromosome divergence between QX1410 and VX34 is similar to that found on the autosomes (one SNV per 131 bp for the autosomes and one SNV per 147 bp for the X chromosome), in contrast to AF16 and QX1410 and both *C. elegans* comparisons, where SNP density is approximately 50% less on the X chromosomes relative to the autosomes (Figure S10). Together, our results add to the growing number of studies showing that the genomes of selfing *Caenorhabditis* species harbor unexpectedly high levels of genetic diversity.

### Selfing *Caenorhabditis* species have undergone more genome rearrangements than their outcrossing sister species

Early comparisons between the *C. elegans* and *C. briggsae* genomes revealed a strikingly high rate of intrachromosomal rearrangement when compared with other taxa such as *Drosophila* (Coghlan and Wolfe *2002; Hillier et al. 2007)*. Intriguingly, the rate of rearrangement was higher than expected from amino acid divergence alone: the rate of amino acid divergence was two times higher than in *Drosophila*, but the rate of rearrangement was four times higher (Coghlan and Wolfe 2002). One possible reason for this discrepancy is that both *C. elegans* and *C. briggsae* reproduce predominantly by self-fertilization. Evolutionary theory predicts that rearrangements that are deleterious in heterozygotes are more likely to become fixed in populations of selfing species than they are in outcrossing species because selfers reach homozygosity more quickly (Lande 1979; Charlesworth 1992). Thus, over time, the genomes of selfing species are expected to undergo more rearrangement than their outcrossing relatives. In *Caenorhabditis*, selfing has evolved three times independently (in *C. elegans, C. briggsae*, and *C. tropicalis*; Figure 4A; (Kiontke et al. 2004, 2011)) and chromosomally resolved reference genomes are now available for *C. inopinata* and *C. nigoni*, the outcrossing sister species of *C. elegans* and *C. briggsae*, respectively (Yin et al. 2018; Kanzaki et al. 2018). Although a chromosomally resolved reference genome was recently published for *C. tropicalis* (Noble et al. 2021), no chromosomally resolved reference genome for its outcrossing sister species, *C. wallacei*, has been published.

**Figure 4:**
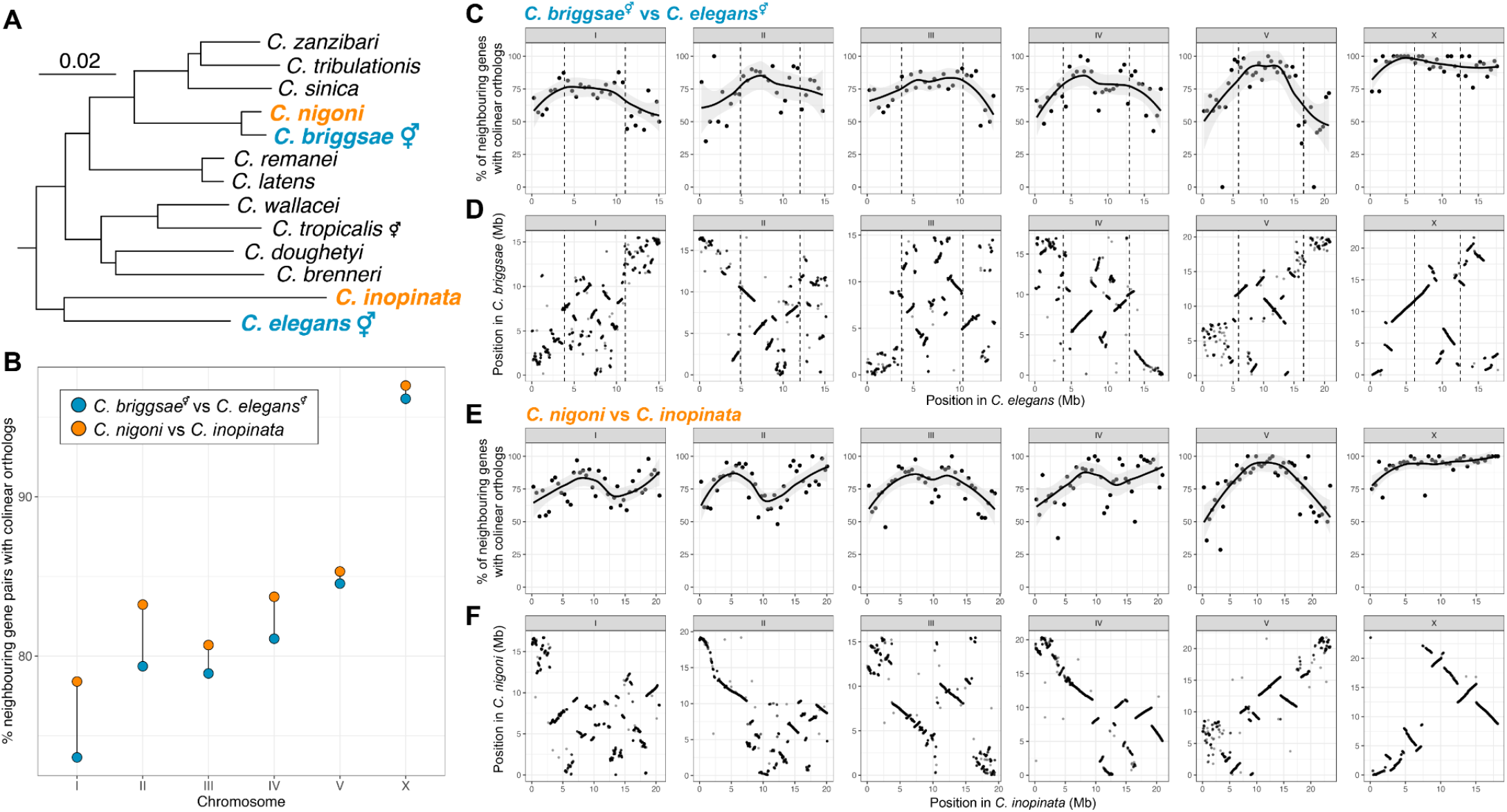
Selfing species have undergone more genome rearrangement than their outcrossing sister species. A) *Caenorhabditis* phylogeny showing relationships within the *Elegans* group (Stevens et al. 2020). The species under comparison are highlighted in blue (selfers) or orange (outcrossers). Branch lengths are in substitutions per site; scale is shown. B) Percentage of neighboring gene pairs in each chromosome with collinear orthologs between the two selfing and two outcrossing species. C) The proportion of neighboring genes in 500 kb windows of the *C. elegans* genome that have collinear orthologs in the *C. briggsae* genome. Solid represent LOESS smoothing functions fitted to the data. Dotted lines represent the positions of the recombination rate domain boundaries (“arms” and “centers”) in *C. elegans* (Rockman and Kruglyak 2009). D) *Positions of 9,461 one-to-one orthologs in the C. elegans* and *C. briggsae* genomes. Dotted lines represent the positions of the recombination rate domain boundaries (“arms” and “centers”) in *C. elegans*. E) The proportion of neighboring genes in 500 kb windows of the *C. inopinata* genome that have collinear orthologs in the *C. nigoni* genome. Lines represent LOESS smoothing functions fitted to the data. F) Positions of 9,461 one-to-one orthologs in the *C. inopinata* and *C. nigoni* genomes.

We sought to determine if the unusually high intrachromosomal rearrangement rate previously observed in *C. elegans* and *C. briggsae* was related to their reproductive mode. As the rate of rearrangement is so high in the genomes of *Caenorhabditis* species, accurately inferring individual rearrangement events is challenging. To circumvent this problem, we identified single-copy orthologs across all four taxa and measured synteny between each genome by counting the number of neighboring gene pairs that had collinear orthologs. Consistent with previous work, we find that genomes of *C. elegans* and *C. briggsae* are highly rearranged relative to one another: an average of 17.7% of neighboring genes in *C. elegans* are non-collinear in *C. briggsae* (Figure 4B) and the autosomal arms are substantially more rearranged than the autosomal centers (Figure 4C, D). Moreover, the X chromosome is substantially more syntenic than the autosomes (96.2% compared with a mean of 79.4%) with the three largest blocks of collinear genes (290, 152, 146) all found on the X chromosome (Figure 4C, D). Consistent with expectations from evolutionary theory, we find that the genomes of *C. elegans* and *C. briggsae* are more highly rearranged than their outcrossing sister species, *C. inopinata* and *C. nigoni* (17.7% of neighbouring genes are rearranged in the selfers compared with 15.3% in the outcrossers; Figure 4E, F). This difference is despite the outcrossing pair of species being more distantly related than the corresponding pair of selfing species (an average of 0.11 amino acid substitutions per site between *C. elegans* and *C. briggsae* compared with 0.13 amino acid substitutions per site between *C. inopinata* and *C. nigoni*; Figure 4A). Although all six chromosomes are less syntenic in selfers than in outcrossers, the difference is not distributed equally. Chromosome I shows the largest reduction in synteny in selfers with approximately 4.8% more rearrangement between neighboring orthologs when compared to the outcrossers. By contrast, chromosome V and the X chromosome, which is hemizygous in males, show comparatively small differences in synteny (0.8% more rearrangements between neighbouring orthologs in selfers than in outcrossers).

However, we note that the patterns we observe could be explained by change or stasis in one of the four species, rather than a general increase in the rearrangement rate in selfing species. To address this point, we compared each species to an independent outcrossing species, *C. remanei*, for which a chromosomally resolved genome assembly was recently published (Teterina et al. 2020). Consistent with our previous results, we find that the genomes of *C. inopinata* and *C. nigoni* are more syntenic (86.1% and 87.7%, respectively) with *C. remanei* than their selfing sister species, *C. elegans* and *C. briggsae* (84.3%, and 86.8%, respectively) (Figure S11). Thus, it appears that the genomes of *C. briggsae* and *C. elegans* species have undergone a higher rate of rearrangement than their outcrossing relatives.

## Discussion

### High-quality reference genomes for two C. briggsae strains

The AF16 reference genome was sequenced nearly 20 years ago using a combination of Sanger-based shotgun sequencing and a physical map generated by sequencing fosmid and bacterial artificial chromosomes (BAC) libraries (Stein et al. 2003). Since that time, advances in sequencing technologies, particularly long-read sequencing and high-throughput mapping approaches, have meant that it is now relatively straightforward to generate reference genomes that far surpass the quality of their predecessors. Despite being one of the highest quality reference genomes available for any eukaryotic organism, long-read sequencing of the *C. elegans* laboratory reference strain recently extended the reference genome by 1.8 Mb, largely by expanding previously collapsed repeated regions, including the rDNA cluster (Tyson et al. 2018; Yoshimura et al. 2019). Our new *C. briggsae* reference genomes, generated using Oxford Nanopore long-read and Hi-C data, are substantially more contiguous and complete than the existing AF16 reference genome. However, despite the quality of our raw data and the relatively small size of the *C. briggsae* genome, some regions of the QX1410 and VX34 genomes remain unresolved, particularly in the subtelomeric regions, which are known to be repetitive (Yoshimura et al. 2019; Kim et al. 2019). It is possible that the base-level accuracy of our long-read data, which we estimate to be approximately 94%, may have prevented large and highly similar repeats from being resolved during assembly. Technologies capable of producing long reads with high base-level accuracy, such as PacBio HiFi sequencing (Wenger et al. 2019), are now available and can lead to substantial improvements in assembly contiguity, largely because of the ability to resolve large, near-identical repeats (Nurk et al. 2020). Future resequencing efforts using these technologies may fully resolve these complex regions of the *C. briggsae* genome.

Despite substantial differences in experimental design, our estimates of the recombination rate in the *C. briggsae* genome and the physical position of the rate boundaries are largely congruent with previous estimates (Hillier et al. 2007; Ross et al. 2011). One region where we notice a large difference is the inferred physical position of the left arm-center boundary on chromosome II, which was estimated to be at approximately 3.4 Mb in our analysis but 4.53 Mb in the analysis of Ross *et al*. 2011. In our recombination map, the region surrounding the boundary shows a gentle change in recombination rate, leading to a 5- to 10-fold increase in standard error of the boundary position relative to other arm-center boundaries. With the exception of the left arm of I and right arms of II and III, our within-domain estimates of recombination rates are similar to previous measurements. The linear segment fitted to the left arm of chromosome I may be skewed because of the lack of a resolved subtelomeric region in the reference genome. In chromosome II, the disparity in marker density between the left and right arms prevented proper estimation of the left arm-center boundary. Subsequently, the skew in the fitted segment may explain the massive increase in recombination rate of the chromosome II right arm relative to other arm regions and to previous measurements.

### Improved approaches to automated protein-coding gene prediction

Complete and accurate gene models are an essential resource to study the biology of any organism. Because of the complexity of protein-coding genes in eukaryotes, all currently available gene prediction tools fail to resolve the structure of most genes correctly (Mathé et al. 2002). For example, the widely used gene prediction pipeline BRAKER only predicts 55% of the *C. elegans* genes accurately (Hoff et al. 2016). Recent advances in long-read RNA sequencing technologies now enable the sequencing of full-length transcripts that serve as an ideal template to accurately infer gene structures ([CSL STYLE ERROR: reference with no printed form.]; Amarasinghe et al. 2020). Using the *C. elegans* N2 reference genes as a truth set, we developed a gene prediction pipeline that effectively combines long- and short-read data and leads to substantial improvements in the sensitivity and BUSCO completeness relative to gene sets predicted using either dataset alone. Based on protein sequence length similarity to the *C. elegans* proteome, we show that our QX1410 and VX34 gene models generated using this pipeline have improved accuracy relative to AF16 gene models generated using other automated approaches (WS255 release, personal communication with WormBase staff). However, our benchmarks on *C. elegans* also reveal thousands of gene predictions that disagree with curated reference models, suggesting that prediction errors remain a common problem, even with long-read data. Manual inspection of gene models and underlying transcriptome data often reveals correctable mistakes such as the retention of non-coding sequences, incomplete coding sequences, missing or additional exons, and fused or split genes. Recent manual curation efforts in *Pristionchus pacificus* and *Haemonchus contortus* have led to substantial improvements in the quality of the reference annotation for these two species (Rödelsperger 2021; Doyle et al. 2020). Manual curation using multiple sources of experimental evidence will be necessary to further improve the quality of the QX1410 genome annotation.

### Hyper-divergent regions punctuate the genomes of wild C. briggsae isolates

The availability of multiple reference genomes for *C. briggsae* allowed us to investigate the pattern of divergence across the genome. *C. briggsae* is known to harbor higher levels of genetic diversity than *C. elegans*, which appears to be explained by the existence of several, well defined phylogeographic groups (Cutter et al. 2006; Félix et al. 2013). Indeed, we found that the genome-wide divergence between QX1410, a member of the “Tropical” group, and VX34, a divergent strain isolated from China, was higher than that between the *C. elegans* laboratory strain N2 and the XZ1516, the most divergent *C. elegans* strain currently known (Lee et al. 2021). A previous population genomic study identified two *C. briggsae* strains isolated in Kerala, India (JU1341 and JU1348) that were far more divergent than any others, including VX34 (Thomas et al. 2015), and thus our data suggest that within-species divergence in *C. briggsae* is substantially higher than is currently known for *C. elegans*. Moreover, the X chromosome in the Keralan strains shows higher levels of divergence than the autosomes, a pattern that is mirrored in the outcrossing sister species, *C. nigoni* (Thomas et al. 2015). High-quality genomes for these strains may provide important insights into the mechanisms and patterns of genomic divergence that preceded speciation.

The presence of hyper-divergent haplotypes in *C. briggsae*, even between the “Tropical” QX1410 and AF16 strains, means that the regions are now known to exist in all three selfing *Caenorhabditis* species (Thompson et al. 2015; Lee et al. 2021; Noble et al. 2021). Although their origins remain unclear, the prevailing hypothesis is that these regions represent remnants of genetic diversity present in the outcrossing ancestor that have been maintained by long-term balancing selection since the evolution of selfing (Thompson et al. 2015; Lee et al. 2021). In *C elegans*, these haplotypes are enriched for genes involved in environmental responses and include the large chemosensory G-protein-coupled receptors (GPCRs) and nuclear hormone receptors (NHRs) (Thompson et al. 2015; Lee et al. 2021). Intriguingly, the structure of the *C. elegans* genome, with large rapidly evolving gene families being enriched on autosomal arms, is known to be conserved in *C. briggsae* (Hillier et al. 2007) raising the possibility that balancing selection has acted on loci with similar functions, or even orthologs, in both species. Furthermore, the presence of these haplotypes in *C. briggsae* also provides an opportunity to conclusively identify their origins. Unlike in *C. elegans*, which lacks a closely related sister species (Kanzaki et al. 2018), *C. briggsae* is estimated to have diverged from its outcrossing sister species, *C. nigoni*, around 3.5 million years ago (Thomas et al. 2015). Future species-wide genome sequencing efforts in *C. briggsae* and *C. nigoni*, especially those involving high-quality genomes of wild isolates generated using long-read data, could identify whether hyper-divergent haplotypes are shared between these two species and whether their divergence is consistent with long-term balancing selection or recent adaptive introgression.

### Selfing species have undergone more genome rearrangement than their outcrossing sister species

One of the first comparative genomic analyses performed between *C. elegans* and *C. briggsae*, two species that have independently evolved self-fertilization from an outcrossing ancestor, was to compare synteny, or gene order, between their genomes, which revealed an unusually high rate of intrachromosomal rearrangement (Coghlan and Wolfe 2002; Hillier et al. 2007). Using chromosomally resolved reference genomes for five *Caenorhabditis* species (C. elegans Sequencing Consortium 1998; Kanzaki et al. 2018; Yin et al. 2018; Teterina et al. 2020), we have shown that the genomes of *C. elegans* and *C. briggsae* have more highly rearranged genomes than their outcrossing sister species, *C. inopinata* and *C. nigoni*. This finding is consistent with theoretical predictions that structural rearrangements that are deleterious when heterozygous will be more likely to become fixed in highly selfing populations because they reach homozygosity more quickly (Lande 1979; Charlesworth 1992). The pattern of reduced synteny in selfers is true across all chromosomes but is less pronounced on chromosomes V and X. A potential explanation for the X chromosome difference is that, in an outcrossing *Caenorhabditis* species, the X is only heterozygous in females (males are hemizygous for X), meaning the effect of selfing on the rearrangement rate in the X chromosome would be expected to be half of that observed in autosomes (assuming a 50/50 ratio of males and females). However, the X chromosome shows a far reduced rearrangement rate than the autosomes in both outcrossers and selfers, suggesting gene order on the X is under selective constraints that are unrelated to reproductive mode. We also note that the overall decrease in degree of synteny seen in selfing genomes is relatively small, which is expected based on the relatively short time each species is believed to have been reproducing via self-fertilization (within the last 3.5 and 4 million years for *C. briggsae* and *C. elegans*, respectively (Cutter et al. 2008; Thomas et al. 2015)). Therefore, although the rates inferred by Coghlan and Wolfe (2003) were likely biased by comparing two selfing species, it remains true that *Caenorhabditis* genomes undergo a high rate of intrachromosomal rearrangement. Because we only surveyed two selfing and outcrossing species pairs, it remains possible that the differences we observed are the result of coincidental lineage-specific variation in rearrangement rate. Therefore, it will be interesting to know if the pattern holds true in the remaining selfing and outcrossing *Caenorhabditis* species pair, *C. tropicalis* and *C. wallacei*, and in related genera that have independently evolved self-fertilization, such as *Oscheius* (Baïlle et al. 2008).

### Future outlook

Although an improved reference genome for *C. briggsae* is an essential step in our efforts to understand genome evolution in *Caenorhabditis*, it is only a beginning. In recent years, an improved understanding of the natural ecology of *Caenorhabditis* nematodes has led to a dramatic increase in the discovery of new species with over 60 species now in laboratory culture, and multiple isolates are available for many species (Kiontke et al. 2011; Félix et al. 2014; Kanzaki et al. 2018; Ferrari et al. 2017; Stevens et al. 2019; Crombie et al. 2019; Dayi et al. 2021). Although some species have high-quality reference genomes (C. elegans Sequencing Consortium 1998; Kanzaki et al. 2018; Yin et al. 2018; Noble et al. 2021; Teterina et al. 2020), the majority have been sequenced using short-reads only and therefore have highly fragmented genome assemblies that obscure the higher order structure of the genome and complicate downstream analyses (Stevens et al. 2019). Efforts to generate chromosomally resolved reference genomes from across the *Caenorhabditis* phylogeny and in related genera will help to answer several long-standing questions about *Caenorhabditis* and nematode genome evolution. Why do genes rarely move between chromosomes despite strikingly high rates of within-chromosomal rearrangement? What is the origin of the ‘arms’ and ‘centers’ recombination landscape, and how well conserved is it? And why, despite nematode chromosomes being holocentric, is the *C. elegans* karyotype so highly conserved in related species? Moreover, in the majority of cases, only a single strain has been sequenced for each species, and studies of within-species genetic variation have been largely restricted to selfing species. Resequencing datasets, particularly those involving long-read data, will reveal the distribution and levels of genetic diversity in outcrossing species and provide an important context to the recent discoveries of hyper-divergent haplotypes in selfers. These new insights, made possible by advances in sequencing technology, an improved understanding of ecology, and intense sampling efforts, will help to place *C. elegans* and the vast body of knowledge of its biology within a rich evolutionary context.

## Methods

### Nematode culture

Nematodes were reared at 20°C using *Escherichia coli* OP50 bacteria grown on modified nematode growth medium (NGMA), containing 1% agar and 0.7% agarose to prevent animals from burrowing (Andersen et al. 2014).

### DNA extraction

To extract DNA, we transferred nematodes from three 10 cm NGMA plates spotted with OP50 E. coli into a 15 ml conical tube by washing with 10 mL of M9. We then used gravity to settle animals on the bottom of the conical tube, removed the supernatant, and added 10 mL of fresh M9. We repeated this wash method three times allowing the animals to settle for 45 minutes each time to serially dilute the *E. coli* in the M9 and allow the animals time to purge ingested *E. coli*. Genomic DNA was isolated from 50 to 300 µl nematode pellets using the Blood and Tissue DNA isolation kit (cat# 69506, QIAGEN, Valencia, CA) following established protocols (Cook et al. 2016). The DNA concentration was determined for each sample using the Qubit dsDNA Broad Range Assay Kit (cat# Q32850, Invitrogen, Carlsbad, CA).

### Short-read Illumina sequencing

To extract DNA, we transferred nematodes from three recently starved 10 cm NGMA plates originally spotted with OP50 *E. coli* into a 15 ml conical tube by washing with 10 mL of M9. We then used gravity to settle animals on the bottom of the conical tube, removed the supernatant, and added 10 mL of fresh M9. We repeated this wash method three times to serially dilute the *E. coli* in the M9 and allow the animals time to purge ingested *E. coli*. Genomic DNA was isolated from 100 to 300 µl nematode pellets using the Blood and Tissue DNA isolation kit (cat# 69506, QIAGEN, Valencia, CA) following established protocols (Cook et al. 2016). The DNA concentration was determined for each sample using the Qubit dsDNA Broad Range Assay Kit (cat# Q32850, Invitrogen, Carlsbad, CA).

For high-coverage sequencing, libraries were generated with New England BioLabs NEBNext® Ultra™ II FS DNA Library Prep (NEB, Ipswich, MA). Samples were sequenced at the Duke Center for Genomic and Computational Biology, Novogene, or the Northwestern Sequencing facility, NUSeq. All samples were sequenced on the NovaSeq 6000 platform (paired-end 150 bp reads).

For low-coverage sequencing, libraries were generated using a modified Illumina Nextera Sample Prep (Illumina, FC-121-1030) protocol. For each sample, 0.16 ng of DNA was tagmented for five minutes at 55°C with 2.5 µl of a 1/35 dilution of the Illumina Transposome in a tris buffer (10 mM Tris-HCl, pH 8.0; 5 mM MgCl2). Tagmented samples were then amplified and barcoded using ExTaq (TaKaRa, RR001B) and custom primers. Resulting libraries were pooled and sequenced on the Illumina MiSeq.

### Long-read Oxford Nanopore sequencing

Nematodes were collected using the same technique as for short-read sequencing but on 14 10 cm plates instead of three. Animals were transferred from plates using 25 ml of M9 and washed into a 50 ml conical. The 300 to 500 µl worm pellets were submitted to the DNA Technologies and Expression Analysis Cores at University of California, Davis for High Molecular Weight gDNA extraction, library preparation, and sequencing on the Oxford Nanopore PromethION system.

### Hi-C library preparation

The Hi-C libraries were prepared using a modified protocol based on a method described previously (Crane et al. 2015). Briefly, approximately 12,000 adult nematodes were harvested and washed in M9 buffer. The animals were cross-linked with 2% (v/v) formaldehyde solution, then dounced to disrupt pellets in 1 ml lysis buffer (10mM Tris-HCl, pH=8.0, 10mM NaCl and 0.1%(v/v) protease inhibitors). The chromatin was digested overnight by *Dpn*II, then incubated at 65°C for 15 minutes to deactivate the enzyme. The DNA ends were biotinylated at 23°C for four hours and blunt-end ligated with T4 DNA ligase at 16°C for four hours. Proteinase K (50 µl of 10 mg/ml) was added to each tube to reverse crosslinks and degrade proteins. Then, equal volumes of phenol and chloroform (1:1) were added per tube and DNA purified using 15 ml phase lock tubes (1500 g for 5 minutes). Then, the aqueous phase was transferred to the clear 35 ml tube and precipitated with 10% volume of 3 M sodium acetate and 2.5 volumes of ice cold 100% ethanol. The pellet was dried at room temperature and then dissolved in 5 mL of TE buffer. Next, biotin was removed from the unligated ends by adding 5 µl of T4 DNA polymerase (NEB) to 5 ug of DNA, before shearing the DNA to a size of 100-300 bp using the Coveris M220 apparatus. Biotinylated fragments were pulled down using streptavidin beads and resuspended in a ligation buffer. Then, Illumina indices were added along with adapters. The beads were pelleted using a Magnetic Particle Concentrator (Thermofisher) for 1 min, and washed several times before being resuspended the beads in 20 µl NEBuffer 2 (New England Biolabs). The final library was generated using the Illumina TruSeq kit (Illumina) and sequenced using the Illumina HiSeq 4000 platform, yielding 50 bp paired-end reads

### RNA extraction

Three sets of samples were collected for *C. briggsae* strains QX1410 and VX34 and *C. elegans* PD1074 in order to maximize transcript representation. For each strain, we sampled a mixed-staged population prepared by chunking plates every two days for several generations, a male-enriched population created by setting up crosses and expanding the population for 2-3 generations, and a starved population by allowing plates to starve out so that they contained dauers and arrested L1 and L2 larvae. For these cultures, nematodes were reared as elsewhere, except that standard nematode growth medium was used (no agarose). Each sample was collected from one 10 cm plate in 100 ml S Basal, flash frozen in liquid nitrogen, and stored at -80°C. RNA was extracted using 1 mL of the TRIzol reagent (Invitrogen, catalog no. 15596026) following the manufacturer’s protocol except that 1 mL of acid-washed sand (Sigma, catalog no. 27439) was added to aid homogenization. RNA was resuspended in nuclease-free water. A Nanodrop spectrophotometer (ThermoFisher) was used to assess the purity of the extracted RNA, a Bioanalyzer (Agilent) was used to determine RNA integrity with the RNA 6000 Pico Kit (Agilent, catalog no. 5067-1513), and a Qubit (ThermoFisher) was used to determine RNA concentration with the Qubit RNA HS Assay kit (ThermoFisher, catalog no. Q32852). Following QC, we pooled 1.5 mg RNA from each sample (mixed-stage, male-enriched, and starved) and used the RNeasy MinElute Cleanup kit (Qiagen, catalog no. 74204) to further purify and concentrate the pooled RNA. We eluted the RNA in nuclease-free water and performed another round of QC as before.

### Long-read RNA sequencing

300 ng total RNA was used to prepare each PacBio Iso-Seq full-length transcript sequencing library. Libraries were prepared in the Duke Center for Genomic and Computational Biology’s Sequencing and Genomic Technologies Core Facility using the NEBNext Single Cell/Low Input cDNA Synthesis and Amplification Module (NEB, catalog no. E6421) and SMRTbell Express Template Prep Kit 2.0 (Pacific Biosciences, catalog no. 100-938-900). Each library was sequenced with 3 SMRT cells.

### Short-read RNA sequencing

We prepared Illumina RNA-seq libraries of *C. briggsae* strains QX1410 and VX34 in one 96-well plate simultaneously. For each sample, the NEBNext Poly(A) mRNA Magnetic Isolation Module (New England Biolabs, catalog no. E7490L) was used to purify and enrich mRNA from 1 µg of total RNA. We performed RNA fragmentation, first and second strand cDNA synthesis, and end-repair processing using the NEBNext Ultra II RNA Library Prep with Sample Purification Beads (New England Biolabs, catalog no. E7775L). Adapters and unique dual indexes in the NEBNext Multiplex Oligos for Illumina (New England Biolabs, catalog no. E6440L) were used to adapter-ligate the cDNA libraries. We performed all procedures according to manufacturer protocols. We determined the concentration of each RNA-seq library using Qubit dsDNA BR Assay Kit (Invitrogen, catalog no. Q32853). RNA-seq libraries were pooled and qualified with the 2100 Bioanalyzer (Agilent) at Novogene, CA, USA. The pooled libraries were sequenced on a single lane of an Illumina NovaSeq 6000 platform, yielding 150-bp paired-end (PE150) reads.

### RIL construction, genotyping, and cross object creation

Recombinant inbred lines between QX1410 and VX34 were constructed by first generating heterozygous individuals by crossing males of each parent strain to hermaphrodites of the other strain. These heterozygous individuals had maternal contributions (mitochondria) from either parent, so four different genotypes were generated: males and hermaphrodites each with either QX1410 or VX34 mitochondria. Heterozygous males from each parental cross were crossed to both types of heterozygous hermaphrodites in four different crosses. Twenty-five hermaphrodite cross progeny from each of these four crosses were picked to individual plates for a total of 100 independent recombinant progeny. These individuals were selfed by single-animal passage for ten generations. After which time, each RIL was cryopreserved and its genome was sequenced.

Once sequenced, raw reads for the 99 lines were processed for genotyping using the Andersen Lab’s nil-ril nextflow pipeline (https://github.com/AndersenLab/nil-ril-nf). The raw reads were aligned to the QX1410 reference genome. Strains that were run multiple times were merged into a single BAM file. Variants were called and put into a dataset along with the parental genotypes. A hidden-markov-model (HMM) was used to fill in missing genotypes from the low-coverage data. The HMM VCF was then used to generate a genotype coordinate flat file for the position of each variant and the parental genotype of each strain at each position.

### Genome Assembly

We downsampled the ONT long read data for QX1410 and VX34 to ∼200x coverage using FiltLong (v0.2.0; https://github.com/rrwick/Filtlong), based on a genome size of 106 Mb. We assembled the subsampled long reads independently for each strain using Canu r10117 (Koren et al. 2017), Flye v2.8.1-b1676 (Kolmogorov et al. 2018) and wtdbg2 v0.0 (Ruan and Li 2019) using default parameters. We used nucmer v3.1 (Delcher et al. 2003) to align each assembly to the current version of the AF16 genome. The Flye assemblies were consistently and substantially more contiguous than the Canu and wtdbg2 assemblies and were broadly congruent with the AF16 reference. We used the ONT-specific error correction tool, Medaka v1.1.2 (https://github.com/nanoporetech/medaka), to correct sequencing errors in the Flye assemblies. Prior to running medaka, we aligned the ONT reads to the Flye assemblies using minimap2 v2.17-r941 (Li 2018) and provided the resulting alignments to racon v1.4.13 (Vaser et al. 2017) to perform error correction using the parameters recommended by ONT (-m 8 -x -6 -g -8 -w 500). We then provided this corrected assemblies to Medaka along with the ONT reads. We corrected remaining sequencing errors by aligning a paired-end Illumina dataset to the assemblies using bwa mem v0.7.17-r1188 (Li 2013) and providing the resulting alignments to Pilon v1.23 (Walker et al. 2014) (using the --fix bases parameter).

### Hi-C scaffolding

To scaffold the polished Flye assemblies into complete chromosomes, we downsampled the Hi-C data to ∼50x coverage using seqtk v1.3-r106 (https://github.com/lh3/seqtk). We used the Juicer/3D-DNA pipeline v1.6 and v180114 (Dudchenko et al. 2017; Durand et al. 2016) to align the Hi-C data to the assembly and scaffold into chromosomal scaffolds (using default parameters, with ‘*Dpn*II’ as the restriction enzyme). We noticed that 3D-DNA was erroneously breaking contigs in highly repetitive regions, and that this behavior could not be suppressed by turning off misjoin correction (-r 0). To avoid this, we used the early exit mode in 3D-DNA (--early-exit) and then manually edited the assembly file to create large chromosomal scaffolds based on the corresponding Hi-C contact map. The resulting assemblies both comprised six chromosomal scaffolds. The QX1410 genome had five unplaced scaffolds (15-71 kb; 183 kb in total span) and the VX34 genome had three unplaced contigs (37-66 kb; 141 kb in total span).

### Mitochondrial genome

We identified the mitochondrial contig by searching the genome assembly with BLASTN using the AF16 mitochondrial genome as a query. As is common for circular genomes, the assembled contig consisted of a “concatemer” composed of more than one copy. To resolve this step, we used mitoHiFi (v1; https://github.com/marcelauliano/MitoHiFi) to decircularize the mitochondrial genome.

### Curation and QC

We assessed the base-level accuracy of the Hi-C-scaffolded assembly by estimating QV score using Merqury v1.1 (Rhie et al. 2020) and the Illumina short-read library. For the unplaced contigs, we manually inspected each contig and the corresponding reads using gap5 (Bonfield and Whitwham 2010). We also aligned each contig to the assembly using BLASTN. We found that all unplaced contigs comprised redundant repeats, several of which were low complexity (including a contig that was composed entirely of repeat units of the telomeric hexamer, TTAGGC). All unplaced contigs were removed from the final assembly. The resulting, curated chromosomes were reoriented relative to the *C. briggsae* AF16 reference genome.

### Repeat Masking

Prior to gene prediction, we masked repetitive sequences using a custom repeat library. The approach used to generate custom repeat libraries for nematode genomes has been previously described (Berriman et al. 2018; Teterina et al. 2020). In summary, we first identified repetitive sequences *de novo* using RepeatModeler from RepeatMasker v2.0.1 (Smit et al. 2015). We identified transposable elements using TransposonPSI ([CSL STYLE ERROR: reference with no printed form.]). We identified long terminal repeat (LTR) retrotransposons using LTRharvest and LTRdigest from Genome Tools v1.6.1 (Gremme et al. 2013; Ellinghaus et al. 2008). Putative LTR retrotransposon sequences were identified with the *gt-ltrharvest* tool, followed by annotation with the *gt-ltrdigest* tool using HMM profiles from Gypsy Database v2.0 (Llorens et al. 2011), and Pfam domains (Finn et al. 2014) selected from tables SB1 and SB2 of (Steinbiss et al. 2009). Output sequences from *gt-ltrdigest* were filtered using the *gt-select* tool to remove repeat candidates without conserved protein domains. Additionally, we retrieved *C. elegans* ancestral repetitive sequences and Rhabditida-specific repeats from RepBase (Bao et al. 2015) and Dfam (Hubley et al. 2016), respectively. Newly generated and retrieved repeat libraries were merged into a single redundant repeat library. Repeats in the merged library were clustered using VSEARCH v2.14.2 (Rognes et al. 2016) and classified with the RepeatClassifier tool from RepeatModeler. We removed unclassified repeats with BLAST (Camacho et al. 2009) hits to *C. elegans* proteins and soft-masked the genome assemblies using RepeatMasker.

### Protein-coding gene prediction

We generated protein-coding gene predictions using BRAKER v2.1.6 (Hoff et al. 2019). In summary, we aligned short RNA-seq reads for each strain to their respective soft-masked genome using STAR v2.7.3a (Dobin et al. 2013) in two-pass mode with a maximum intron size of 10 kilobases. We then supplied sequence alignments and soft-masked genome assemblies to the BRAKER pipeline. Additionally, we generated high-quality transcripts from Pacific Biosciences (PacBio) long RNA reads using the isoseq3 pipeline v3.4.0 ([CSL STYLE ERROR: reference with no printed form.]). We aligned PacBio high-quality transcripts for each strain to their respective genome using minimap2 (Li 2018). We supplied long-read transcript alignments to StringTie v2.1.2 (Kovaka et al. 2019) and performed transcriptome assembly. The coding sequences of the assembled transcripts were predicted using Transdecoder v5.5.0 ([CSL STYLE ERROR: reference with no printed form.]). We extracted StringTie gene models with incomplete coding sequences (CDS) using *agat_sp_remove_incomplete_gene_models* script from AGAT v0.6.0 (Dainat et al. 2020). We identified StringTie models with incomplete CDS that mapped to loci where no BRAKER gene model was present. We merged BRAKER models with complete StringTie models and incomplete StringTie models that map to new loci using *agat_sp_merge_annotations*.*pl* script from AGAT. We resolved the format of genes that overlap in a single locus using *agat_sp_fix_overlaping_genes*.*pl* from AGAT.

### Gene prediction pipeline benchmarks

We benchmarked the quality of our gene prediction pipeline using the *C. elegans* N2 reference annotation from WormBase (WS279) as a truth set. We generated gene models for the *C. elegans* reference genome using the gene prediction pipeline described previously. For short-read based gene prediction, we generated an N2 mixed-population RNA-seq sample by pooling Illumina data from individual samples representing all larval stages and adults (retrieved from SRR953117, SRR953118, SRR953119, SRR953120, and SRR953121). To match the depth of our *C. briggsae* RNA-seq samples, we randomly downsampled the pooled N2 Illumina reads to 35 million reads. We compared individual BRAKER and StringTie models and merged gene models against the existing *C. elegans* reference annotation using GffCompare from GffRead (Pertea and Pertea 2020). We extracted sensitivity and precision statistics for base, exon, intron, intron chain, transcript, and locus level. We assessed the biological completeness of the N2 predictions using BUSCO v4.0.5 (Simão et al. 2015) with the Nematoda (odb10) database.

### C. briggsae gene predictions

We generated gene models for *C. briggsae* QX1410 and VX34 using the gene prediction pipeline described above. To assess the quality of the *C. briggsae* gene predictions, we developed a script to search for reciprocal BLAST hits between protein sequences of *C. briggsae* predictions and the *C. elegans* reference annotation. We extracted protein sequences from each gene prediction set using GffRead. For each protein sequence, we kept the best BLAST hit based on expected value and bit score. Two or more best hits were kept if their expected values and bit score were identical. We considered a pair of protein sequences to be reciprocal if the best hit of the first sequence matched the best hit of the second, and vice versa. Pair of sequences with multiple best hits were considered reciprocal if any of their hits were reciprocal. For each set of gene predictions, we counted the number of genes that have at least one reciprocal protein sequence (total matches). We compared the protein lengths of every predicted protein sequence against its reciprocal. We counted the number of genes that had the same protein length (1:1 hits) and the number of genes that were within 5% of the protein length (5% off hits) of its reciprocal. We assessed biological completeness using BUSCO with the Nematoda (odb10) database.

### Genetic map construction and domain analysis

We filtered markers and estimated genetic distances using the *R/qtl* package (Broman et al. 2003). In summary, we read RIL genotypes into a cross object using the *read*.*cross()* function with the ‘riself’ parameter. We identified and removed 25 markers with distorted segregation patterns using the *geno*.*table()* function, and three markers that fell outside six major linkage groups using the *formLinkageGroups()* function. Additionally, we developed an algorithm to search and remove individual or groups of markers with local deviations in allele frequency equal or greater than 4% relative to its neighboring markers. We removed 103 markers with deviations in allele frequency. We estimated genetic distances using *est*.*rf()* with the default Haldane map function. We manually removed 22 additional markers with exceedingly high recombination rate in close physical distance and re-estimated genetic distances. Chromosomal domain analyses were performed using the *R/segmented* package (Muggeo 2003). Arm-center domain boundaries were defined using segmented linear regressions with two expected breakpoints. Because markers in the left arm of chromosome II were sparse, the left arm-center boundary was not defined as a segment breakpoint when including all markers in the initial linear fit. To estimate this missing arm-center boundary, we excluded markers after 15 Mb in the right end of chromosome II prior to linear fit. Tip domains were manually called by identifying the first pair of markers that showed a change in recombination rate.

### Assessing genome-wide divergence between the three C. briggsae genomes

To measure and visualise nucleotide divergence between the assemblies, we used nucmer 3.1 (Delcher et al. 2003) to align VX34 and AF16 to the QX1410 reference genome. To measure amino acid divergence, we employed an orthology clustering approach. Briefly, we identified and selected the longest isoform of each protein-coding gene for QX1410, AF16, and VX34 using AGAT v0.5.0 (Dainat et al. 2020). The isoform-filtered proteomes were then clustered into orthologous groups using OrthoFinder v2.5.2 ((Emms and Kelly 2019). We selected all single-copy orthologous groups and aligned the protein sequenced with MAFFT v7.475 (Katoh and Standley 2013) and calculated the average amino acid identity using a custom script (available at https://github.com/lstevens17/briggsae_reference_genome_MS). We also called SNVs using an assembly-based approach. We aligned AF16 and VX34 to the QX1410 reference genome using minimap2 2.17-r941 (Li 2018) and provided the resulting PAF file to paftools v2.18-r1015 (available at https://github.com/lh3/minimap2) to call variants (setting the minimum alignment length to compute coverage and call variants to 1000). The resulting VCF file was filtered to contain only biallelic SNVs using bcftools v1.13 (Danecek et al. 2014) and SNV densities per chromosomes were calculated using bedtools v2.30.0 (Quinlan and Hall 2010).

### Comparing synteny between selfing and outcrossing species

We downloaded the genomes and annotation files for *C. elegans (C. elegans Sequencing Consortium 1998), C. inopinata* (Kanzaki et al. 2018), and *C. nigoni* (Yin et al. 2018) from WormBase (WS279) (Harris et al. 2020), and for *C. remanei* from NCBI (GCA_010183535.1) (Teterina et al. 2020) and used AGAT (v0.5.1; available at https://github.com/NBISweden/AGAT) to extract the longest isoform from each protein-coding gene. We generated two orthology clustering sets by clustering the isoform-filtered protein sequences with OrthoFinder v2.5.2 (Emms and Kelly 2019): one containing only the selfing/outcrossing species pairs (*C. elegans, C. inopinata, C. briggsae*, and *C. nigoni*) and a second that also included *C. remanei*. For both datasets, we selected orthogroups containing protein sequences that were single-copy across all species and extracted their corresponding coordinates using custom scripts (available at https://github.com/lstevens17/briggsae_reference_genome_MS). To count the proportion of neighboring genes that had collinear orthologs, we assigned a number of each orthologous gene, corresponding to its order along each chromosome, and for each neighboring gene pair determined if their orthologs were collinear (*i*.*e*., had a distance of one). These data were then used to compute values for non-overlapping 500 kb windows, identify blocks of collinear genes, and average synteny for each chromosome. We performed two comparisons (1) a direct comparison between *C. elegans* and *C. briggsae* (selfers), and between *C. inopinata* and *C. nigoni* (outcrossers), and (2) a comparison between all four focal species and *C. remanei*. To compare our measures of synteny with amino acid divergence, we used ETE3 (Huerta-Cepas et al. 2016) to extract branch lengths (in amino acids substitutions per site) from a recently published *Caenorhabditis* phylogeny based on the protein sequences of 1,167 single-copy orthologs (Stevens et al. 2020).

## Data and code availability

Raw sequencing data and genome assembly and annotation files have been archived under the NCBI study accession PRJNA784955. Code associated with the analyses and figures can be found at https://github.com/lstevens17/briggsae_reference_genome_MS.

## Acknowledgements

We would like to thank members of the Andersen lab for comments on this manuscript. Most *Caenorhabditis* research, including this work, would not be possible without Wormbase, so we thank them deeply for their contributions. This work was funded by grants to E.C.A. from NIH (R01 ES029930) and the Human Frontiers Science Program (RGP0001/2019). E.C.A. and L.R.B. were also funded by R01 ES029930. A.J.M.W. and H.N. were funded by R35 GM122502. J.D. was funded by the Howard Hughes Medical Institute.

