## Supplementary Information for "Chromosome-level reference genomes for two strains of *Caenorhabditis briggsae*: an improved platform for comparative genomics"

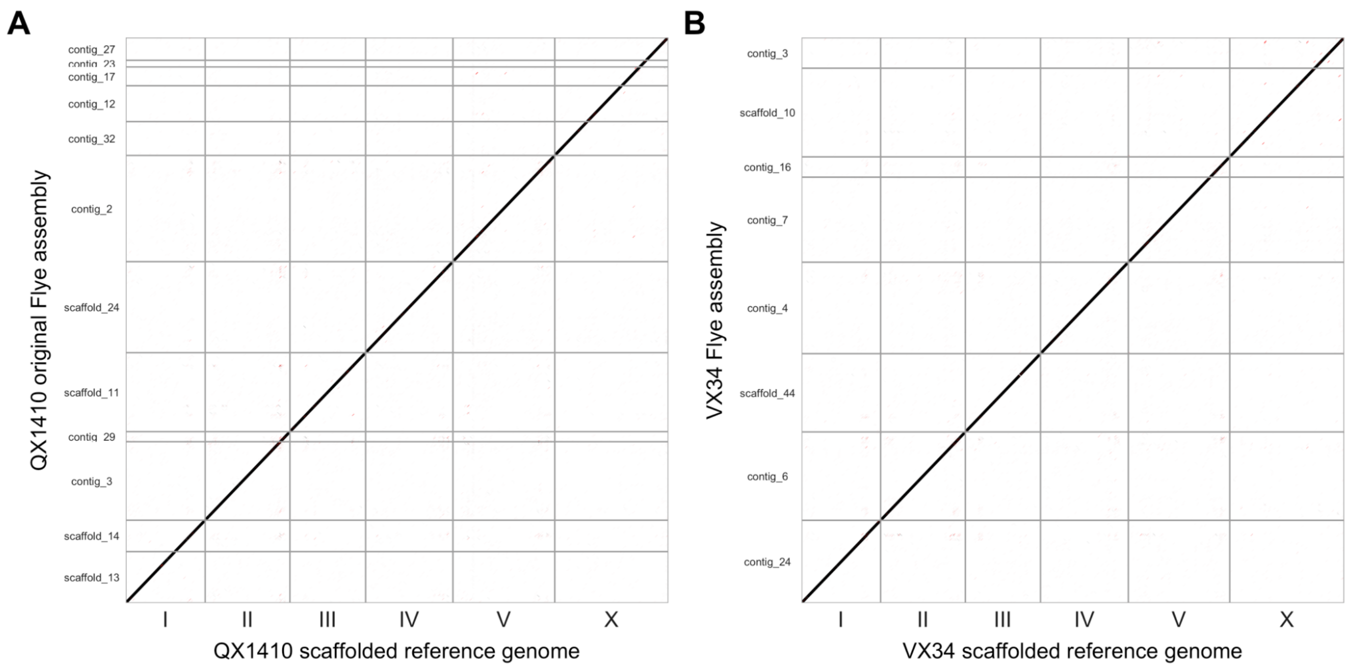


**Figure S1: Highly contiguous assemblies from PromethION long-read data**
Whole-genome alignments between the final scaffolded reference genome and the Flye genome assemblies for (1) QX1410 and (2) VX34. Contigs/scaffolds less than 100 kbp in length are not shown. Three chromosomes were represented by single contigs/scaffolds in the QX1410 Flye assembly (II, IV, V) and four chromosomes were represented by single contigs/scaffolds in the VX34 Flye assembly (I, II, III, IV).

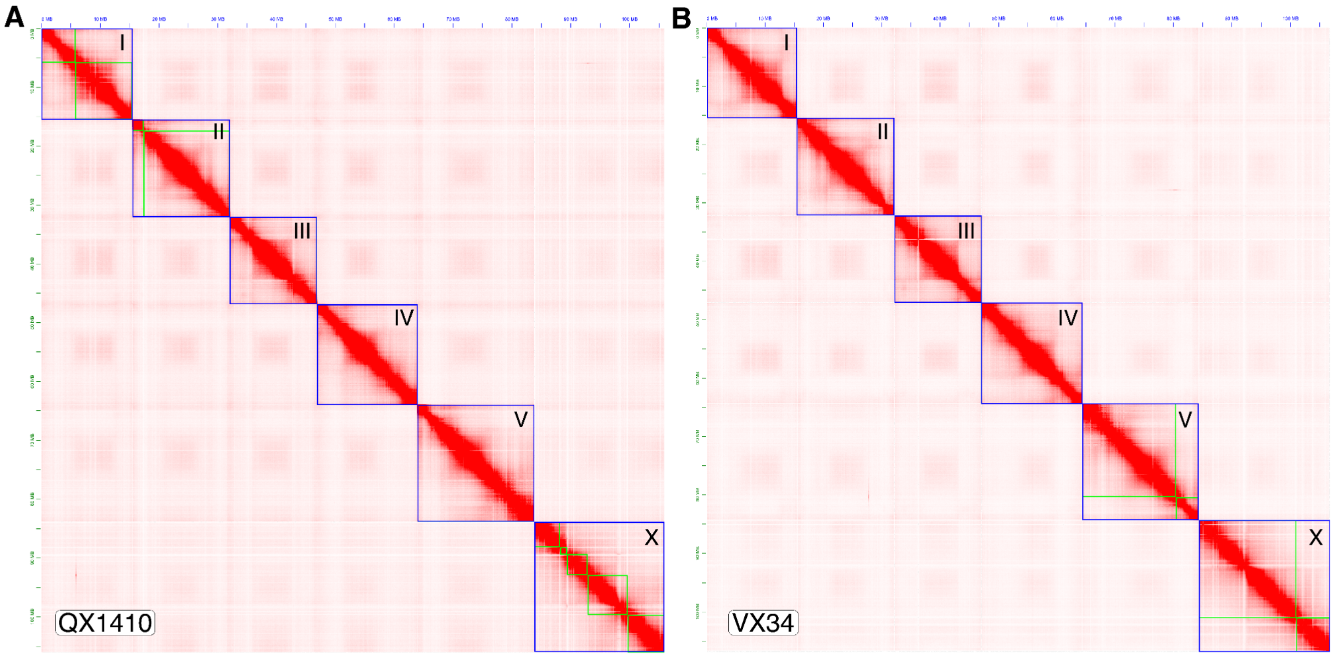


**Figure S2: Hi-C contact maps for QX1410 and VX34.**
Hi-C contact map for (A) QX1410 and (B) VX34. Hi-C read data were mapped to the assembly and processed using the 3D-DNA pipeline. Hi-C maps were visualised using Juicebox. Blue boxes indicate chromosomes; green boxes indicate the Flye contigs or scaffolds that comprise each chromosome. Chromosomes with no green boxes were fully assembled by Flye.

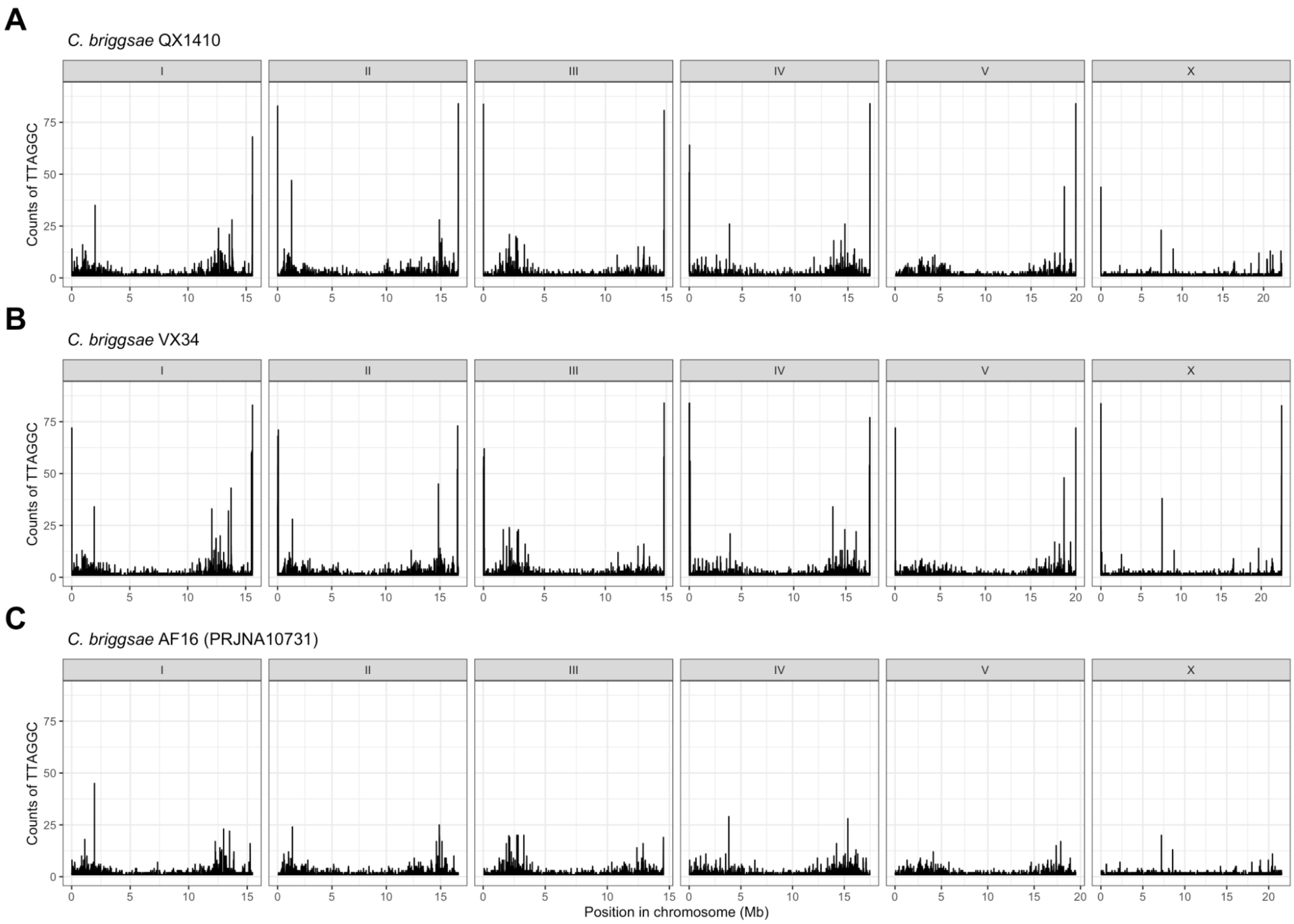


**Figure S3: Telomeric repeat sequence in *C. briggsae* reference genomes**
Counts of the nematode telomeric repeat sequence (TTAGGC) in 1 kb windows in (A) *C. briggsae* QX1410 (B) *C. briggsae* VX34, and C) *C. briggsae* AF16 reference genomes. Telomeric repeats were identified using seqkit (Shen et al. 2016) and binned into 1 kb windows using bedtools (Quinlan and Hall 2010).

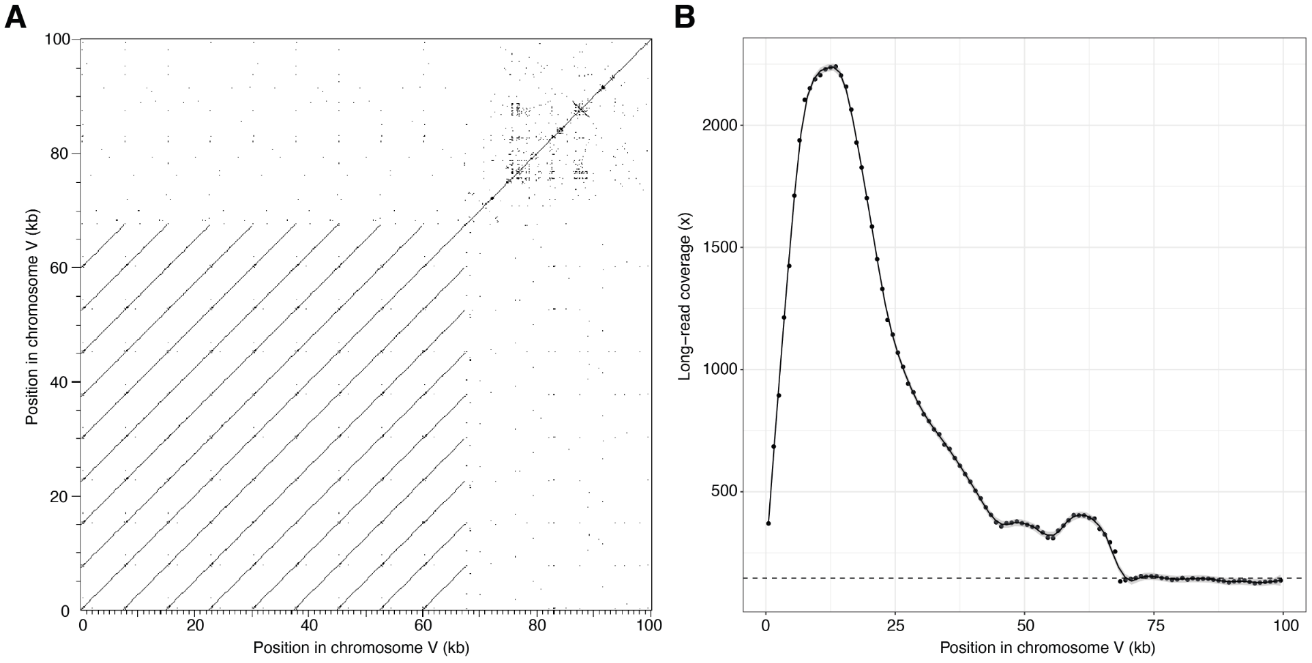


**Figure S4: Collapsed ribosomal DNA cluster in the QX1410 reference genome**
(A) Self-alignment of the first 100 kb of the chromosome V showing the nine repeating ~7.5 kb rDNA units extending to approximately 68 kb. Alignment and plot generated using Dotter.
(B) Coverage of long Oxford Nanopore reads in the first 100 kb of chromosome V. The solid line represents a LOESS smoothing function fitted to the data. The dotted line represents modal coverage for chromosome V (146x). The average coverage for the first 68 kb is 943x, suggesting that the true number of rDNA cistron units is approximately 58 copies.

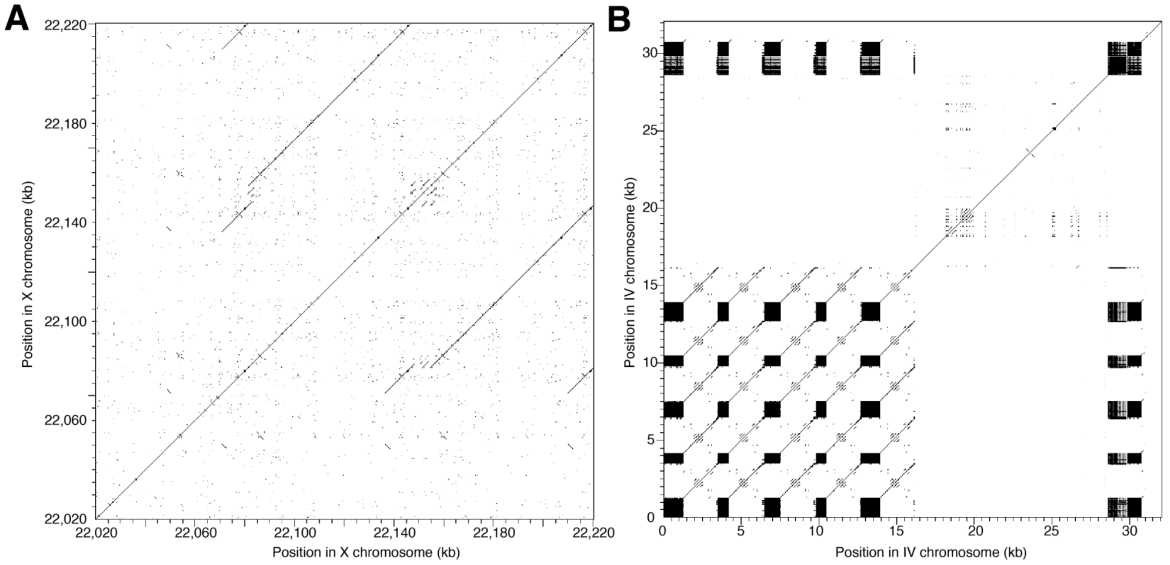


**Figure S5: Highly repetitive subtelomeric regions in the QX1410 reference genome**

(A) Large approximately 65 kb tandem repeat in the right-end subtelomeric region of the X chromosome (22,020-22,220 kb). The right-end of the X chromosome does not terminate in telomeric repeat sequence in our reference genome. Alignments and plots generated using Dotter.

B) Low-complexity repeats in the left-end subtelomeric regions of chromosome IV (0-32 kb). The dark regions comprise blocks of a 19-mer (GGCTTCCCGCTTAGGCTTA) interspersed with blocks of the telomeric repeat sequence (TTAGGC). Alignments and plots generated using Dotter.

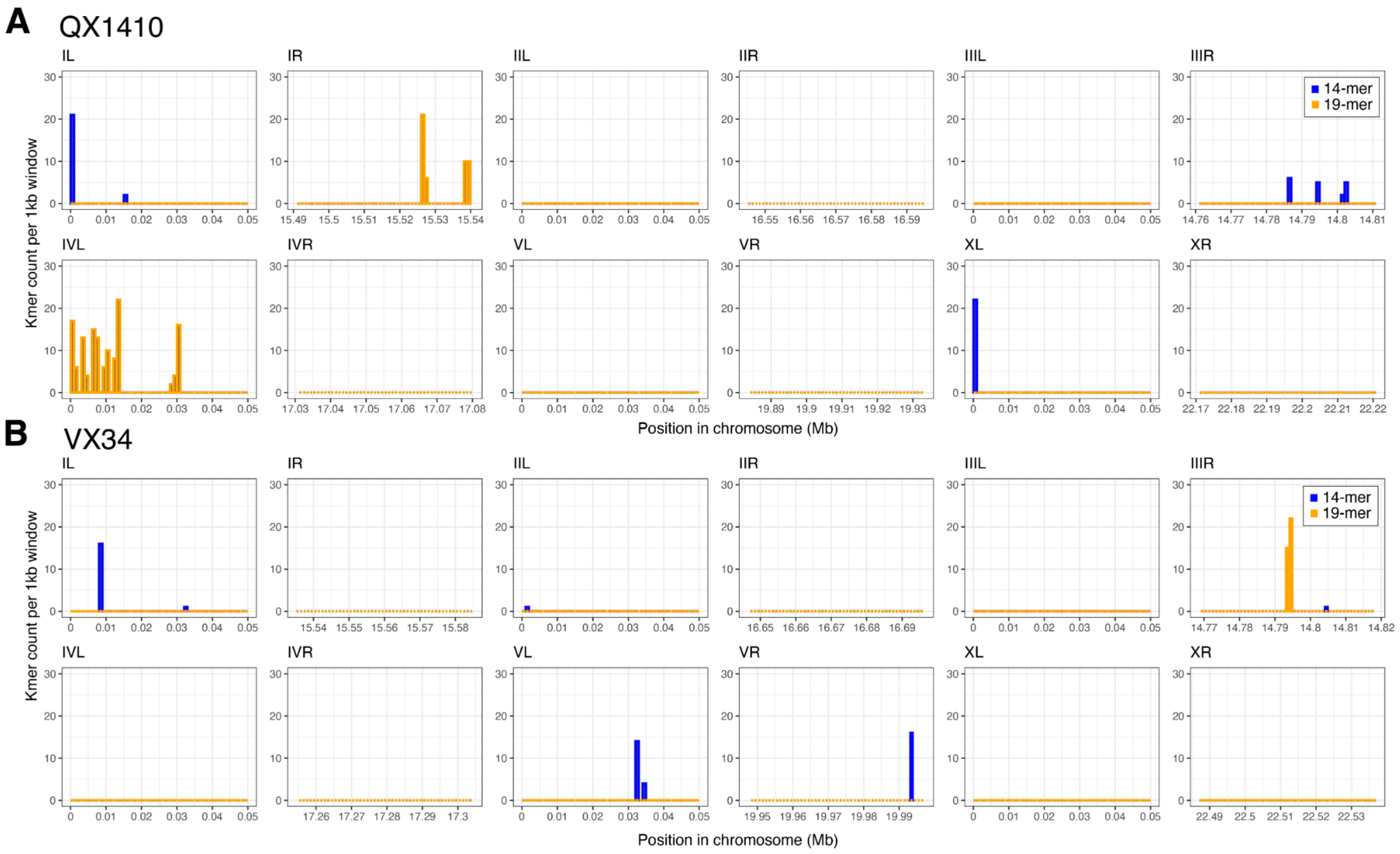


**Figure S6: Blocks of simple repeat in the subtelomeric regions.**
Counts of a 14-mer and 19-mer that are found exclusively at the subtelomeric ends in (A) QX1410 and (B) VX34. Both repeats contain the nematode telomeric sequence (TTAGGC). The first and last 500 kb are shown for each chromosome. Locations of the repeats were found using seqkit locate (v 0.15.0).

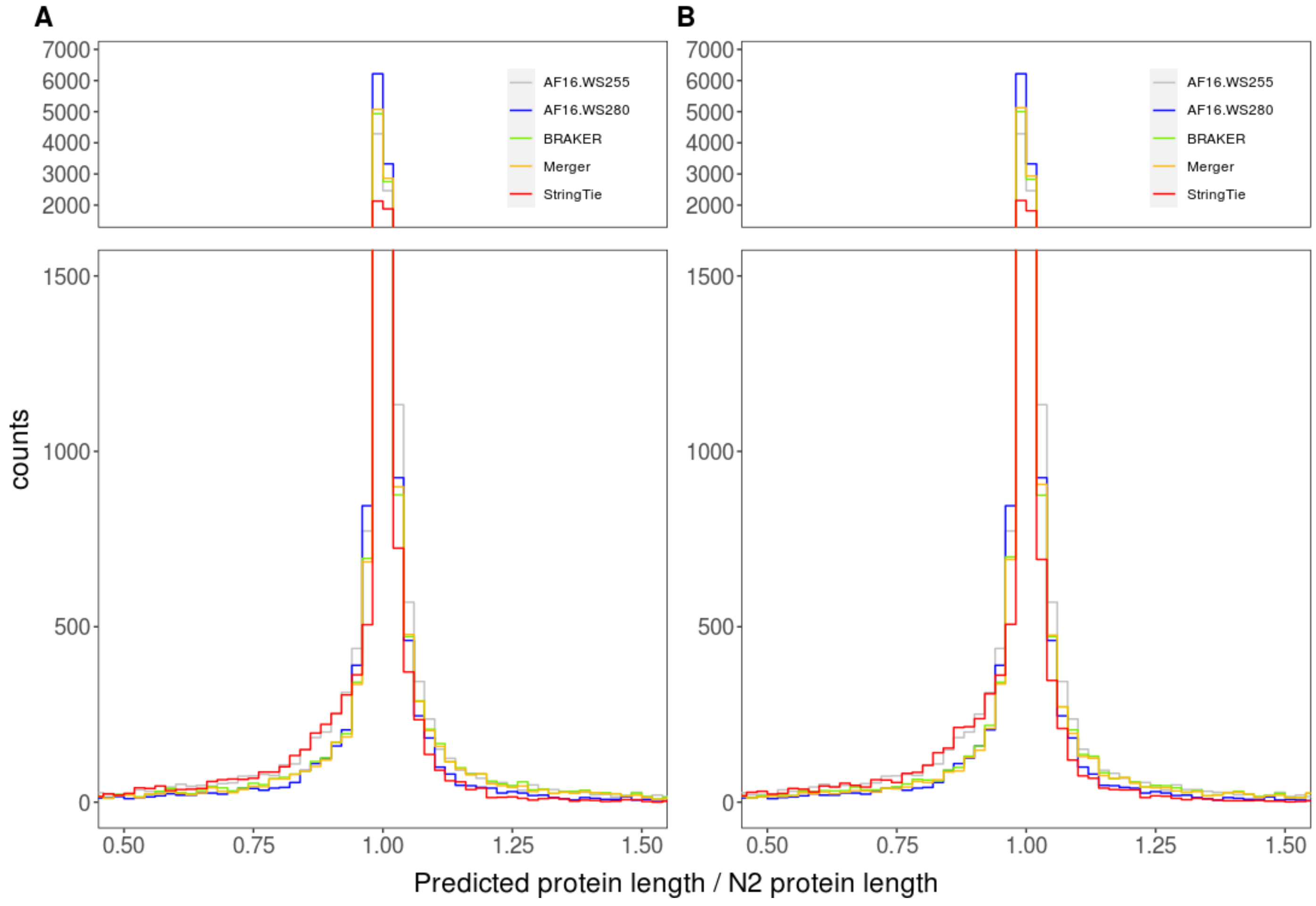


**Figure S7: Protein length accuracy in *C. briggsae* genome annotations.**
Counts of binned protein length ratios calculated from the comparison of the best reciprocal hits between the *C. elegans* reference annotation and genome annotations for *C. briggsae* strains (A) QX1410 and (B) VX34.

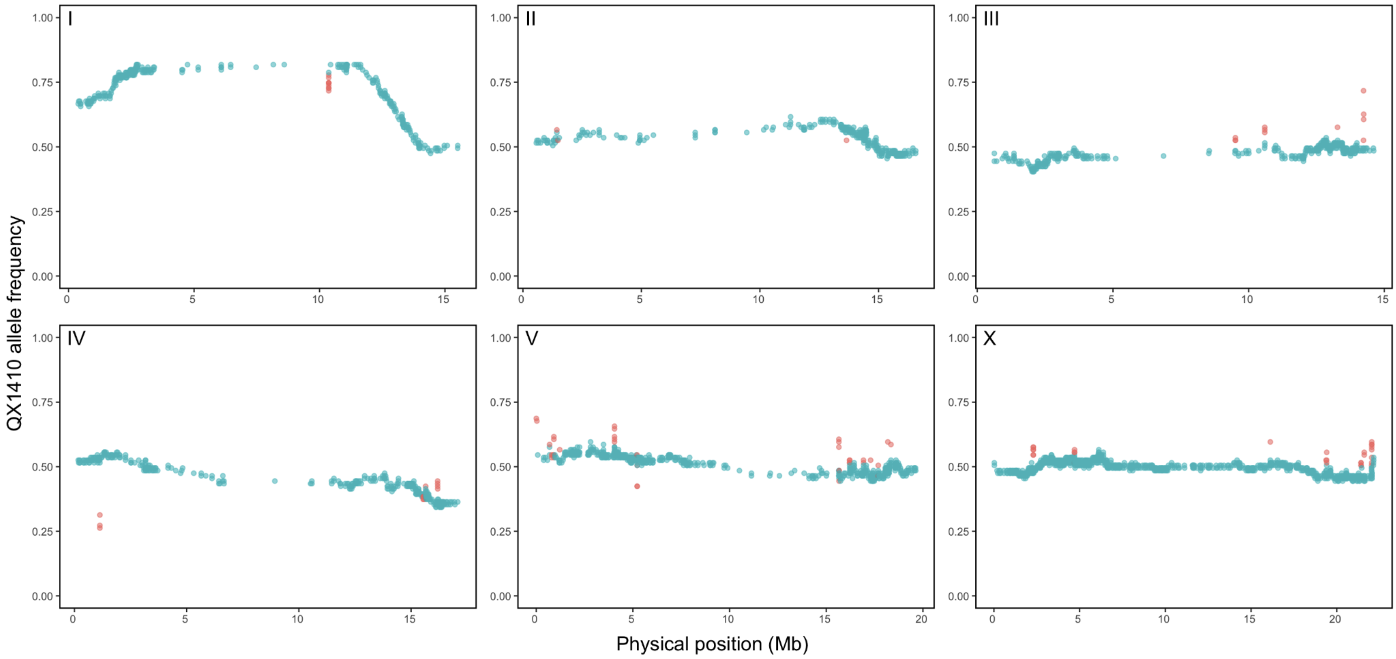


**Figure S8: Markers with local allele frequency deviations.**
Marker allele frequency as a function of physical position. Each dot represents a marker. Red dots are markers that show a deviation in allele frequency higher than 4% relative to its neighboring markers. Deviant markers were removed prior to estimation of the genetic map.

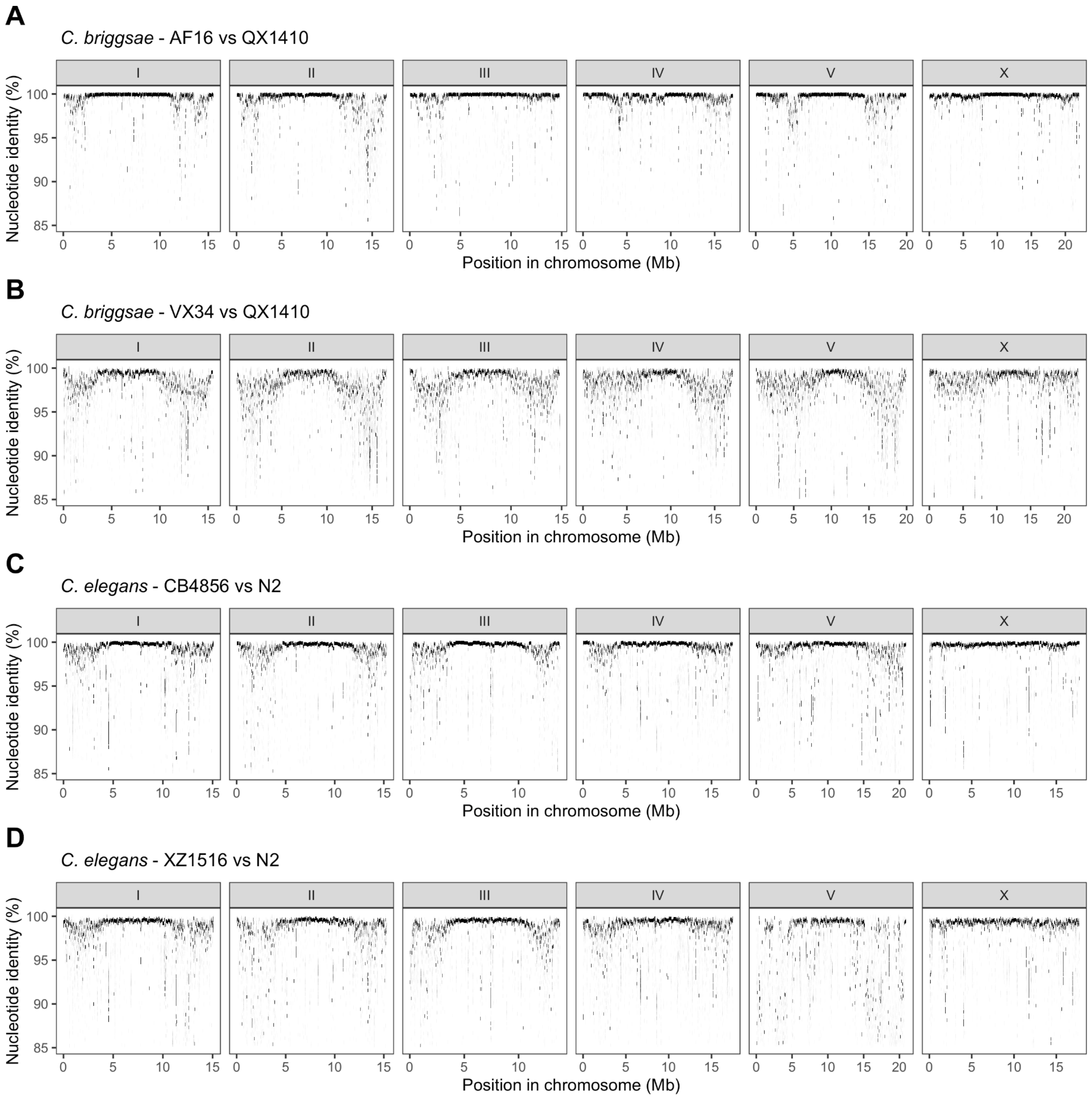


**Figure S9: Genome-wide divergence between three *C. briggsae* and three *C. elegans* strains.** Nucleotide identity between aligned regions of (A) AF16 and QX1410, (B) VX34 and QX1410, (C) CB4856 and N2, and (D) XZ1516 and N2. Genomes were aligned using nucmer and aligned regions of 1 kb or longer are shown.

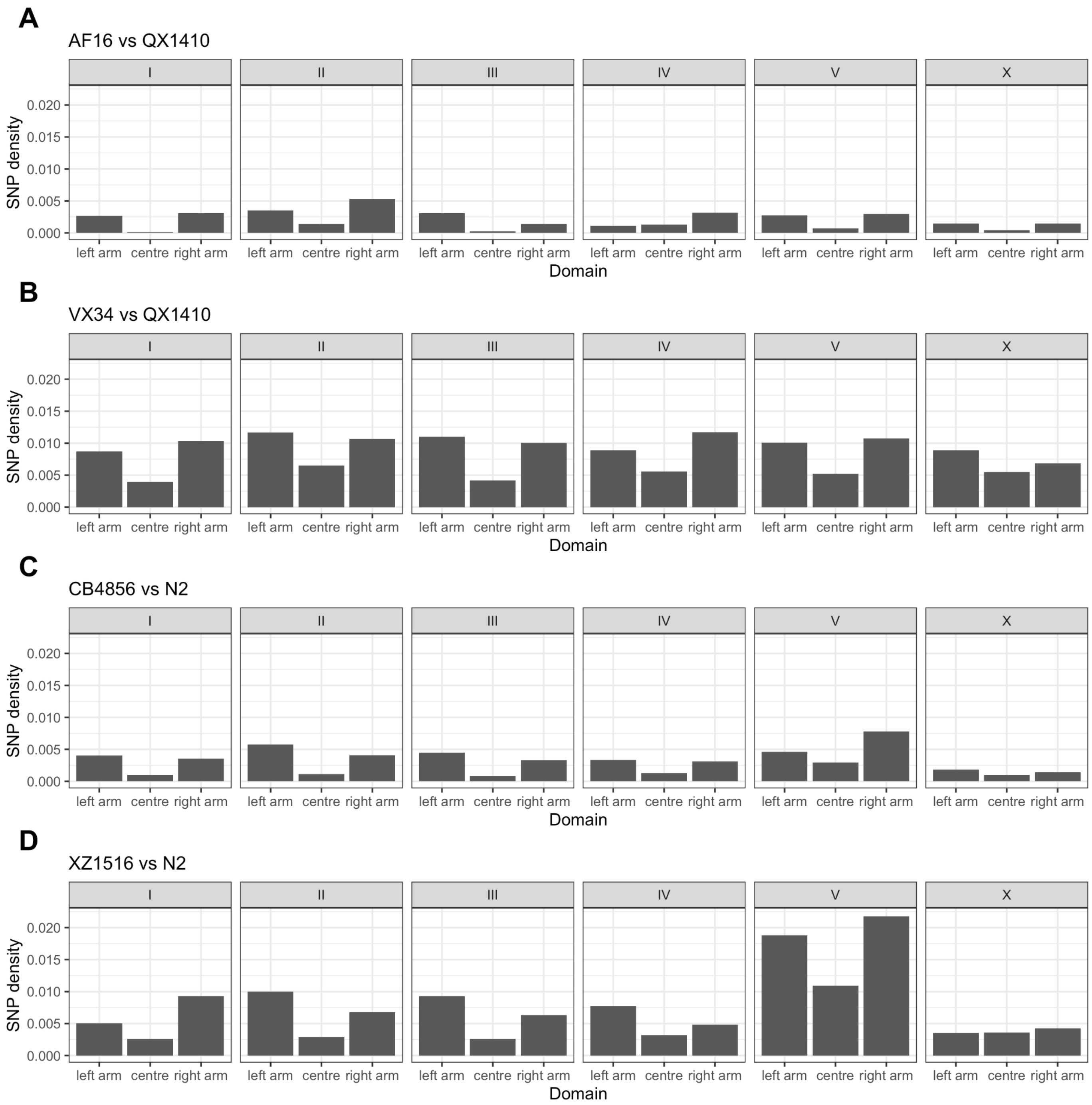


**Figure S10: SNP density by domain in *C. briggsae* and *C elegans***
SNV density in recombination domains between (A) AF16 and QX1410, (B) VX34 and QX1410, (C) CB4856 and N2, and (D) XZ1516 and N2. Variants were called by aligning reference genomes minimap2 and calling variants using paftools. Only biallelic SNVs are shown (indels were filtered from the VCF).

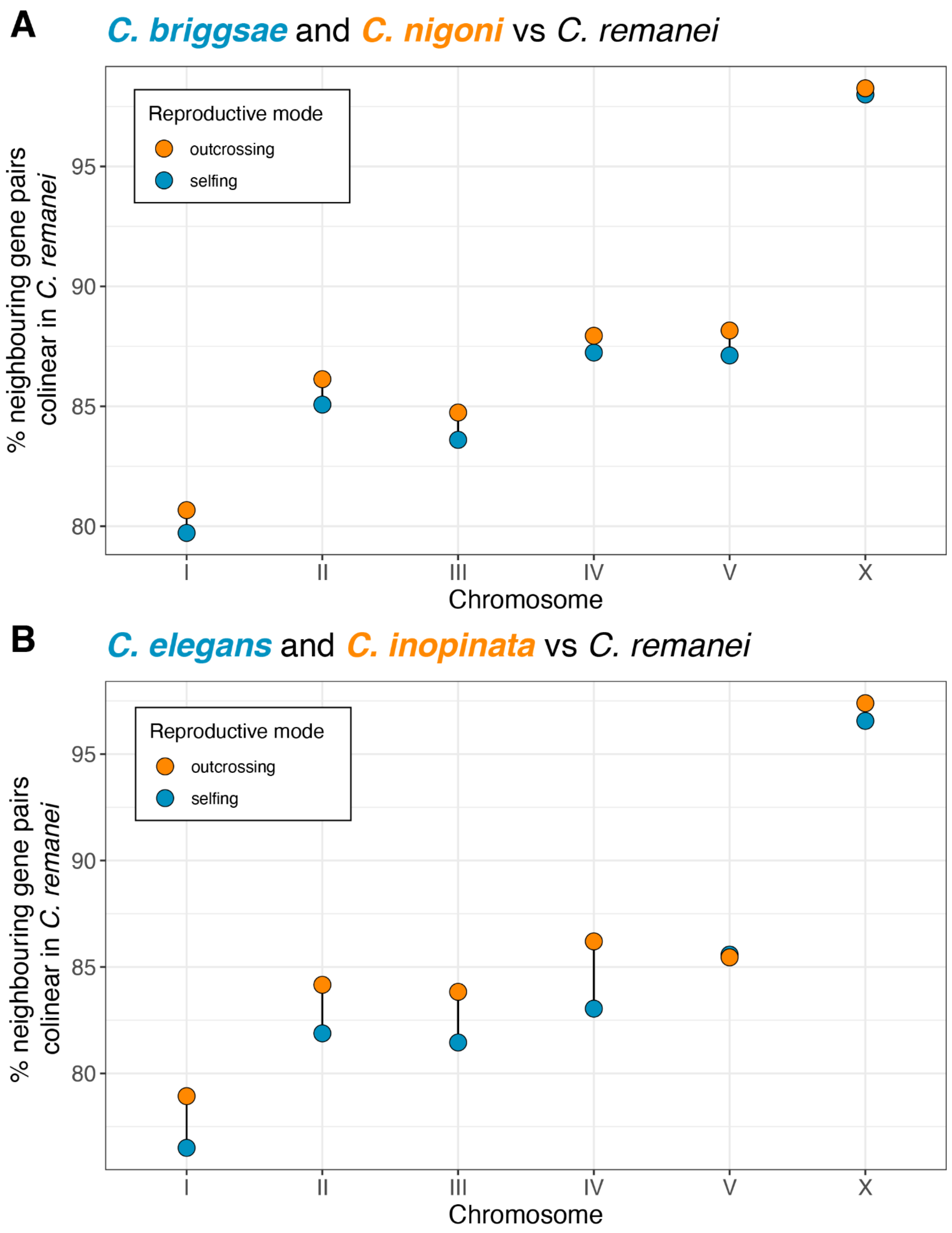


**Figure S11: Selfing species have undergone more genome rearrangement than their outcrossing sister species.** Percentage of neighboring gene pairs in each chromosome with colinear orthologs between the two selfing and two outcrossing species.

**Table S1: Pipeline benchmarks using N2**

| **Sensitivity** | StringTie | BRAKER | Merger (AGAT) |
|---|---|---|---|
| Exon | 48.2 | 68.6 | 74.7 |
| Intron | 54.1 | 84.9 | 86.1 |
| Intron Chain | 23.7 | 38.1 | 43.1 |
| Transcript | 23.0 | 38.0 | 42.9 |
| Gene | 39.3 | 63.5 | 68.2 |
| **Precision** | | | |
| Exon | 90.9 | 75.7 | 75.6 |
| Intron | 96.6 | 87.7 | 87.1 |
| Intron Chain | 64.3 | 52.9 | 45.4 |
| Transcript | 64.3 | 52.8 | 45.4 |
| Gene | 73.8 | 62.4 | 67.9 |
| **Other** | | | |
| BUSCO | 77.9 | 98.6 | 99.1 |
| Matching Transcripts | 7794 | 12890 | 14538 |
| Total Transcripts | 12115 | 24423 | 32000 |
| Total Genes | 10025 | 19135 | 18684 |

**Table S2: Accuracy and completeness of *C. briggsae* reference annotations**

| **Strain** | **Annotation** | **1:1 Matches** | **5% off Matches** | **Total Matches** | **BUSCO (Proteins)** | **Busco (Genome)** |
|---|---|---|---|---|---|---|
| **QX1410** | StringTie | 849 | 5644 | 9218 | 81.2% | 99.4% |
| | BRAKER | 2448 | 9709 | 13809 | 98.6% | 99.4% |
| | Merger | 2490 | 9975 | 13827 | 99.2% | 99.4% |
| **VX34** | StringTie | 858 | 5575 | 9012 | 81.4% | 99.4% |
| | BRAKER | 2481 | 9857 | 13696 | 98.9% | 99.4% |
| | Merger | 2519 | 10113 | 13717 | 99.2% | 99.4% |
| **AF16** | WS255 | 2125 | 9218 | 14189 | 98.1% | 99% |
| | WS280 | 2916 | 11779 | 14689 | 99.2% | 99% |

**Table S3: Summary of removed markers prior to genetic map estimation**

| | |
|---|---|
| Markers showing segregation distortion (automated) | I_8165834, I_8167909, I_9241873, I_9241875, I_9307016, I_10328256, I_10331809, I_10364095, I_10369765, I_10456881, I_10508408, I_10684674, I_11091573, I_11154474, I_11203793, I_11207032, I_11244815, I_11247984, I_11380224, I_11381761, I_11406290, I_11439942, I_11484817, I_11497452, I_11540065 |
| Markers outside six major linkage groups (automated) | I_51741, I_51749, I_51784 |
| Markers with strong local allele frequency deviations (automated) | I_10367053, I_10367090, I_10367167, I_10367188, I_10367222, I_10367260, I_10367329, I_10367420, I_10367483, II_1434153, II_1480993, II_13651562, III_9511353, III_9511927, III_9515020, III_10594337, III_10594372, III_10594487, III_13275435, III_14233798, III_14234616, III_14239231, III_14240335, IV_1132302, IV_1132397, IV_1133178, IV_15536612, IV_15538088, IV_15546661, IV_15551098, IV_15572081, IV_15615967, IV_15641135, IV_15646594, IV_16178922, IV_16179986, IV_16181646, IV_16181712, V_266, V_30227, V_687593, V_787373, V_797378, V_868549, V_906016, V_908194, V_909009, V_909277, V_1212931, V_4046302, V_4046312, V_4046323, V_4046404, V_4046501, V_5214410, V_5217136, V_5218081, V_5218768, V_15652026, V_15652027, V_15652150, V_15659060, V_15670673, V_16205687, V_16207115, V_16211106, V_16211225, V_16924346, V_16926467, V_17292374, V_17688509, V_17689243, V_18183197, V_18346107, X_2298824, X_2299239, X_2299335, X_2299336, X_2299381, X_2299557, X_4701087, X_4702985, X_4703096, X_4709617, X_16124312, X_19377317, X_19412499, X_19412649, X_19414799, X_21410821, X_21411677, X_21420112, X_21427127, X_21603523, X_21603636, X_22030409, X_22039913, X_22039945, X_22039977, X_22040624, X_22040631, X_22041405, X_22042932 |
| Markers with high recombination rate in close physical distance and moderate local allele frequency deviations (manual) | II_12285, II_2216118, II_2218699, II_16571881, II_16571481, II_11779, II_11398, II_11429, III_9518491, III_9517915, III_9518404, III_9516439, IV_49449, IV_50840, IV_46473, IV_49267, V_14255854, V_14255766, V_14255878, V_14256052, X_21407010, X_21406898 |
